# Elevated type I interferon signaling defines the proliferative advantage of ARF and p53 mutant tumor cells

**DOI:** 10.1101/2024.11.11.623046

**Authors:** Alex Mabry, Catherine E. Kuzmicki, Angelina O’Brien, Leonard B. Maggi, Jason D. Weber

## Abstract

The tumor suppressors p53 and ARF collaborate to prevent unwarranted cell proliferation and as such are two of the most frequently mutated genes in human cancer. Concomitant loss of functional p53 and ARF leads to massive gains in cell proliferation and transformation and is often observed in some of the most aggressive human cancer subtypes. These phenotypic gains are preceded by increased type I interferon (IFN) signaling that involves canonical STAT1 activation and a subsequent IFN-stimulated gene (ISG) signature. Here, we show that cells lacking p53 and ARF require active JAK1 to phosphorylate STAT1 on Y701 to maintain their high rate of proliferation. In fact, the use of selective JAK1 inhibitors ruxolitinib or baricitinib inhibited the induction of ISG’s and the proliferation of p53 and ARF deleted cells. We identify a group of solid human tumors that lack functional p53 and ARF, show an expression signature of the upregulated type I IFN response genes, and are sensitive to selective JAK1 inhibitors. These data suggest that the type I IFN response acts as a positive driver of proliferation in the absence of p53 and ARF and, as such, presents itself as a potential therapeutic target in aggressive solid tumors.

## INTRODUCTION

Two of the genes most frequently inactivated in human cancer are the tumor suppressors TP53 and CDKN2A. The p53 tumor suppressor is activated upon DNA damage or cellular stress [1, 2]. Its activation leads to the initiation of transactivated p53 target genes that induce DNA repair, cell cycle arrest, or apoptosis [3, 4]. p53 protein levels are highly regulated, mainly through MDM2, which binds to p53 and targets it for proteasomal degradation [3]. The *CDKN2A* locus (*Ink4a/Arf*) encodes two distinct tumor suppressors, p16INK4a and p19ARF (p14ARF in humans) [6, 7]. P16INK4A has a singular role in binding to CDK4/6 to prevent the activity of the CDK4/6- cyclin D holoenzyme [4, 5]. ARF is canonically known for its p53-dependent tumor suppressive functions [6, 7]. In response to hyperproliferative signals such as those emanating from Ras [8, 9] and Myc [10, 11], ARF is induced [12] and binds to MDM2 [6, 7, 13, 14]. The ARF-MDM2 complex is then sequestered in the nucleolus [15, 16]. This allows nucleoplasmic p53 to perform its transcriptional tumor suppressor activities that lead to cell cycle arrest or apoptosis [15, 17]. However, there is a growing body of work demonstrating that ARF has tumor suppressive functions independent of p53 [18–26]. *In vitro*, ARF overexpression can inhibit *p53*-null cell proliferation [19]. *In vivo*, singly knocked out *p53*-null or *Arf*-null mice develop lymphomas or sarcomas within four months (*p53*-null) or eight months (*Arf*-null) while *p53/Arf*-double null mice exhibit lymphomas or sarcomas in combination with various carcinomas, suggesting that p53 and ARF can independently suppress transformation of epithelial cells [19].

The loss of functional p53 leads to elevated expression of ARF protein in vitro and in vivo, implying that p53 actively suppresses ARF transcription [7, 10, 18, 27, 28]. p53 binds directly to the ARF locus, and then recruits HDACs to deacetylate the locus, which in turn recruits the polycomb repressive complex [28]. These data suggest that ARF is a sensor of p53 fidelity, and this increased ARF can serve as a potential backup tumor suppressor when normal p53 function is lost [18]. This notion is supported by tumor data where co-inactivation of *CDKN2A* and *TP53* is highly prevalent in many human cancers [27–30]. We have previously demonstrated that primary mouse embryo fibroblasts (MEFs) and established human TNBC cell lines with co-inactivation of p53 and ARF have elevated expression of STAT1 and ISG15, and these cell lines are sensitive to STAT1 or ISG15 depletion [18]. Using MEFs deficient in p53, we found that ARF depletion leads to increased IFNβ secretion and STAT1 activation, enhancing the proliferation, transformation, and in vivo tumorigenicity of these cells. Furthermore, we showed *in vitro* and *in vivo* that p53 and ARF collaborate to suppress STAT1 signaling. These data demonstrate that p53 and ARF cooperate to independently suppress an oncogenic IFNβ-STAT1-ISG15 signaling pathway [18].

IFNβ is a type I interferon best known for its role in eliciting an antiviral response. As a primary response to viral infection, cells typically secrete IFNβ [31]. Secreted IFNβ binds to the type I IFN receptor, causing dimerization and activation of Janus kinase 1 (JAK1) and tyrosine kinase 2 (TYK2) [32, 33]. JAK1 and TYK2 activation leads to phosphorylation of STAT1 and STAT2, which can dimerize and lead to transcriptional activation of various interferon-stimulated genes (ISG’s), leading notably to induction of the ISG15 protein [34–36]. Because the type I IFN response during a viral infection typically leads to decreased cellular proliferation and/or apoptosis, the pathway has often been considered tumor suppressive [37]. STAT1 has a variety of tumor suppressive functions through the transactivation of several transcriptional targets, most notably p21 and p53 [40, 41]. STAT1 tumor suppressive functions have also been observed *in vivo*, as *Stat1*-null mice spontaneously form mammary tumors [38, 39]. However, there is an increasing body of evidence suggesting that STAT1 also has tumor-promoting functions. The expression of unphosphorylated STAT1 in tumor cells has been associated with a poor prognosis and can promote the development of sarcomas [44]. In human and mouse breast cancers, STAT1 displays increased expression and activity and contributes significantly to the progression of these cancers [18, 40]. Several other members of the type I IFN pathway have increasing evidence as tumor invasion and tumor cell growth promoters [18, 41–43].

Our previous work demonstrated that loss of p53 and ARF leads to induction of IFNβ, STAT1, and ISG15 and that these components are necessary and sufficient for increased proliferation and tumorigenicity [18]. The mounting evidence that the type I IFN response pathway, and particularly STAT1, contribute to tumorigenesis suggests the need to investigate and characterize the tumor-promoting functions of this pathway. Here, we sought to use *p53*-null MEF lines previously established by Forys et al. [18] to further interrogate the critical components of the type I IFN pathway that drive cell proliferation. We demonstrate that phosphorylation of STAT1 on Y701 by JAK1 is a crucial necessary modification for enhanced proliferation and transformation in cells lacking p53 and ARF. Selective JAK1 inhibitors inhibited STAT1-Y701 phosphorylation, resulting in decreased cell proliferation. Our results point to the potential use of selective JAK1 inhibitors to prevent STAT1 activation and to inhibit the proliferation and transformation of cells lacking the p53 and ARF tumor suppressors.

## RESULTS

### JAK1 is required for the increased proliferation and tumorigenicity of Δp53-Ras-shARF cells

We have previously shown that loss of p53 via Cre-mediated excision of *p53*^fl/fl^ alleles (Δp53) results in dramatic gains in ARF expression [18]. These cells can be transformed by oncogenic *Ras^V12^* alleles, but additional depletion of endogenous ARF using knockdown hairpins specific for ARF (shARF) results in massive gains in proliferation and transformation of these cells (termed Δp53-Ras-shARF cells) [18]. In particular, the increased tumorigenicity of Δp53-Ras-shARF cells is completely dependent on IFNβ production and downstream activation of STAT1 and expression of ISG15 [18]. To further characterize the signaling pathway upregulated in Δp53-Ras-shARF MEFs, we sought to better understand the components responsible for the increased activation of the IFN pathway, using ISG15 expression as a downstream marker of the pathway. In the canonical type I IFN pathway, STAT1 activation occurs through its phosphorylation by Janus kinase 1 (JAK1) [44]. To determine whether JAK1 was necessary for STAT1 activation, we knocked down endogenous JAK1 using two short hairpins in the Δp53-Ras-shARF MEFs. Phospho-STAT1, total STAT1, and ISG15 protein levels were markedly reduced upon JAK1 knockdown (Fig. 1A); however, knockdown of JAK2, which is not involved in the type I IFN response, did not alter ISG15 levels (Supplemental Fig. 1), suggesting that selective activation of JAK1 is responsible for type I IFN pathway signaling in these cells. Furthermore, both short-term (cell counting) and long-term (foci) proliferation was significantly impaired after JAK1 knockdown (Fig. 1B and 1C). The soft agar colonies were also massively reduced in size and number after JAK1 knockdown in Δp53- Ras-shARF MEFs further indicating that the transformation of these cells depends on JAK1 activation (Fig. 1D). Furthermore, this reduction appears to be entirely due to reduced proliferation, since we did not observe any changes in apoptotic markers after JAK1 knockdown (Supplemental Fig. 2). Taken together, these data demonstrate that JAK1 is required for the proliferation and transformation of Δp53-Ras-shARF cells.

**Fig. 1.**
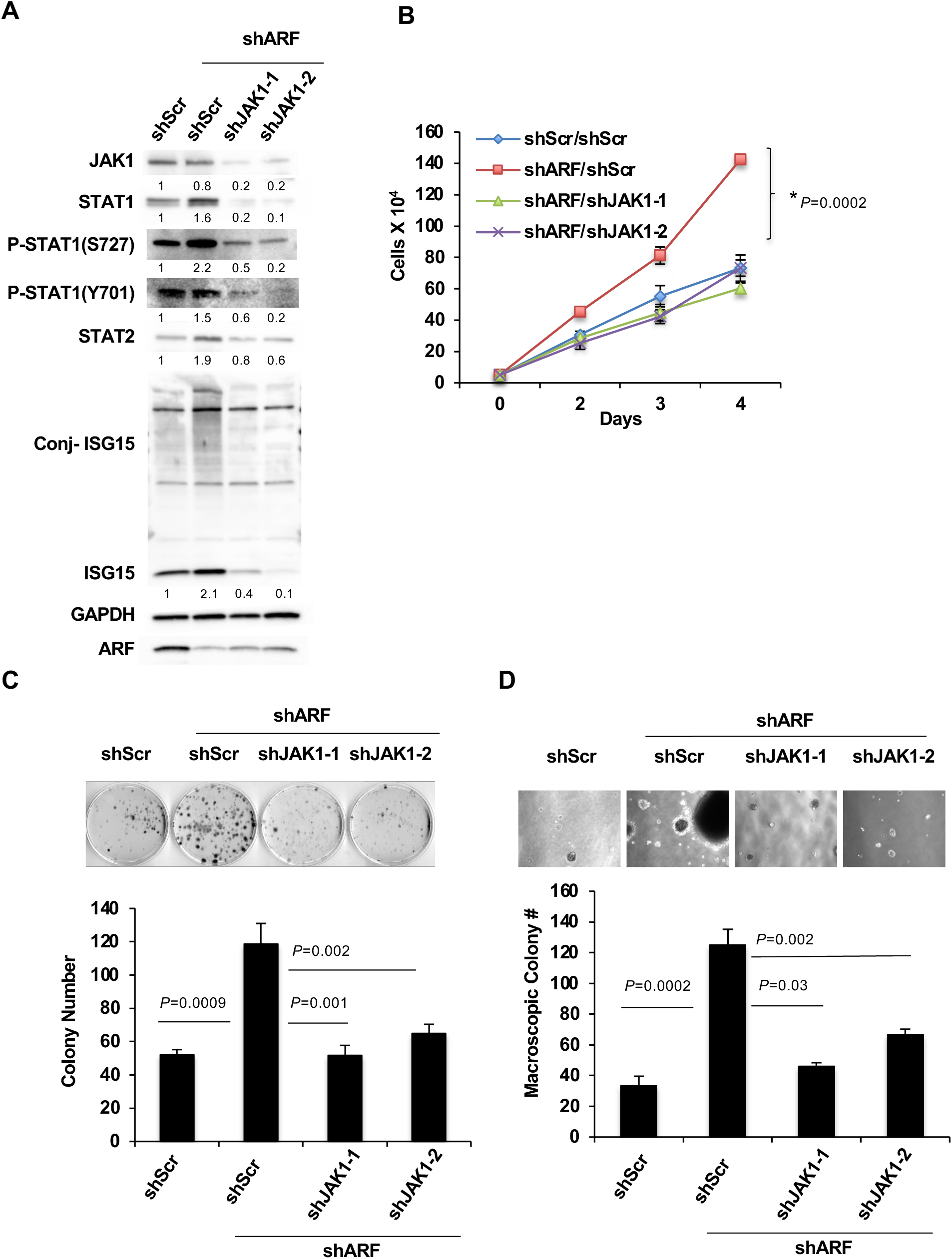
JAK1 is required for increased proliferation and transformation of Δp53-Ras-shARF MEFs. **A** Immunoblot analysis of cell lysates from Δp53-Ras-shScr or –shARF MEFs infected with two specific shRNAs targeting JAK1. Fold levels below each blot are relative to shScr control. Proteins analyzed are provided next to each blot. **B** Four-day proliferation assay performed with cells described in (A). \**P*=0.0002 (green or purple compared to red). **C** Representative image of foci assay performed with cells described in (A). Quantification of three independent measurements is shown (below) with indicated *P* values. Error bars represent standard deviation (SD) of n=3. **D** Representative image of cells described in (A) growing in soft agar. Macroscopic colonies were quantified (below) with indicated *P* values. Error bars represent standard deviation (SD) of n=3.

### The canonical p-STAT1-Y701 site is required for increased proliferation and tumorigenicity of Δp53-Ras-shARF cells

Our data are in stark contrast to the findings that ISG15 transcription can be driven by an unphosphorylated STAT1 complex (U-ISGF3) that forms in response to chronic but low levels of type I IFNs [44]. Δp53-Ras-shARF cells exhibited low levels of secreted IFN-β and the expression of the predicted U-ISGF3-driven genes [18]. We suspected that the phosphorylated form of STAT1 might be driving the ISG15 levels observed in Δp53-Ras-shARF MEFs. JAK-mediated phosphorylation of STAT1 (phospho-STAT1) on Y701 is critical for STAT1 nuclear translocation and downstream transcription of the type I IFN response [45–47]. To determine if phospho-STAT1 was the critical component of the ISGF3 complex, we performed knockdown rescue experiments with a short hairpin targeting endogenous STAT1[18] and retroviral overexpression rescue constructs expressing wild type STAT1 (WT), or individual mutants at two known STAT1 phosphorylation sites (Y701A and S727A)[44–46]. STAT1 suppression led to a dramatic decrease in free and conjugated ISG15 levels (Fig. 2A). This reduction was only rescued with overexpression of STAT1-WT or STAT1-S727A and not with STAT1-Y701F, indicating that the phosphorylation of the Y701 site was critical for STAT1 activity and subsequent downstream transcriptional stimulation, effectively ruling out the U-ISGF3 complex. Only cells with high levels of phospho-STAT1-Y701 and ISG15 retained an increase in short- and long-term proliferation as measured by cell growth and foci assays; respectively (Fig. 2B-D). Again, this was evident in soft agar transformation assays where loss of STAT1 resulted in the failure to form transformed colonies that were only rescued with STAT1 constructs that retained an intact Y701 residue (Fig. 2E and F).

**Fig. 2.**
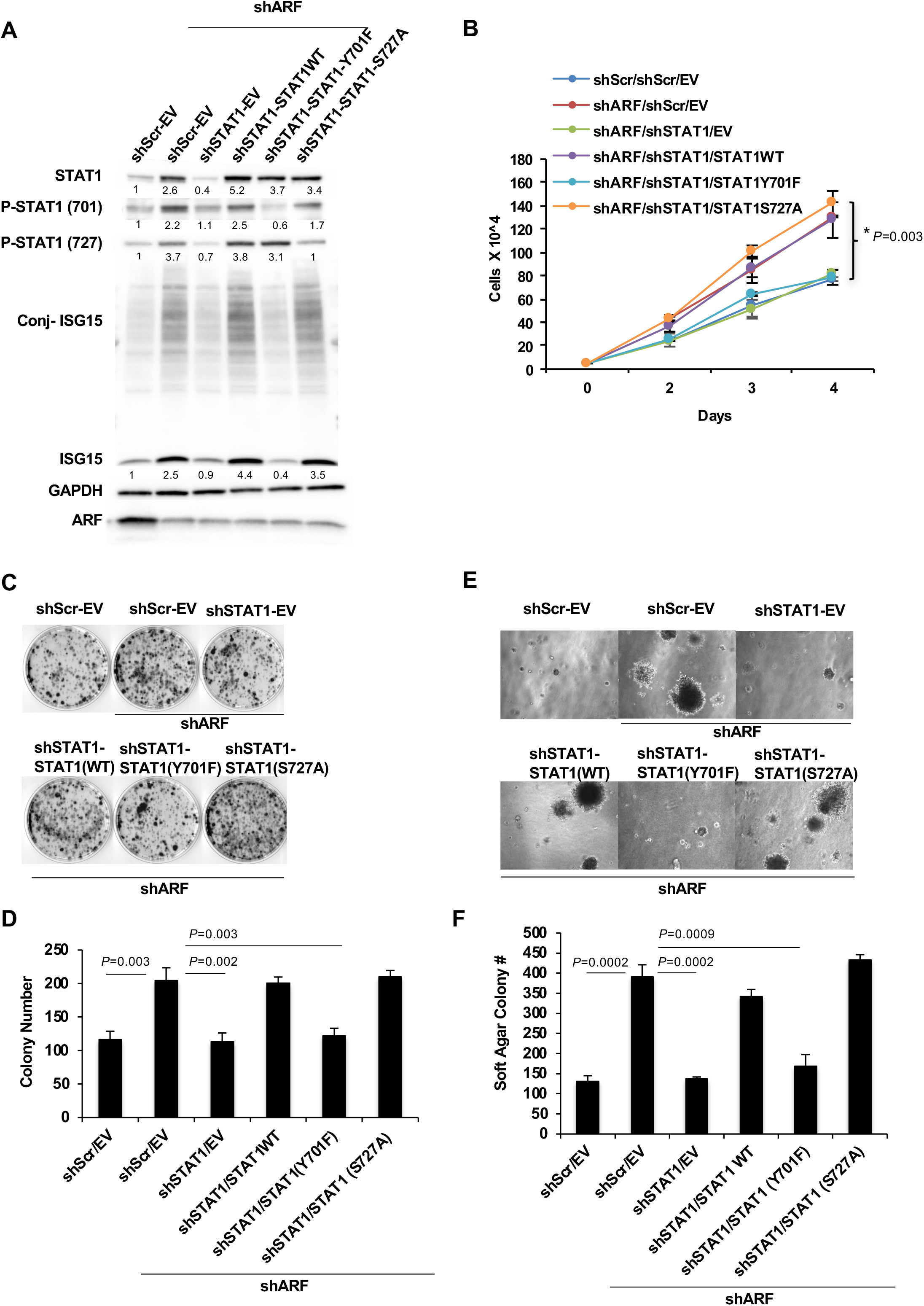
The canonical p-STAT1-Y701 but not p-STAT1-S727 site is required for increased proliferation and transformation of Δp53-Ras-shARF MEFs. **A** Immunoblot analysis of cell lysates from Δp53-Ras-shScr or –shARF MEFs infected with a specific shRNA targeting STAT1 and subsequently infected with retroviral rescue constructs empty vector (EV), STAT1 WT, Y701F, or S727A. Fold levels below each blot are relative to shScr control. **B** Four-day proliferation assay performed with cells described in (A) with indicated *P* values for significant difference only. \**P*=0.003 (light blue compared to purple). **C** Representative images of foci assay performed with cells described in (A). **D** Quantification of three independent measurements is shown with indicated *P* values for significant difference only. Error bars represent standard deviation (SD) of n=3. **E** Representative image of cells described in (a) growing in soft agar. **F** Macroscopic colonies were quantified (below) with indicated *P* values for significant difference only. Error bars represent standard deviation (SD) of n=3.

### STAT2 is required for the increased proliferation and tumorigenicity of Δp53-Ras-shARF cells

Activated STAT1 and STAT2 bind to form a heterodimer that acts as a powerful transcription factor for genes stimulated by interferon [47]. Thus, we sought to determine the impact of STAT2 on STAT1’s ability to induce the observed phenotypes in Δp53-Ras-shARF MEFs. Reduction in STAT2 levels, using short hairpins targeting endogenous STAT2, resulted in significant decreases in ISG15 protein expression (Fig. 3A), loss of proliferation measured by cell growth and foci assays (Fig. 3B-D) and attenuated soft agar colony formation (Fig. 3E and F), similar to that observed for STAT1 loss. ISG15 can be stimulated through numerous STAT and IRF complexes. The dependency on phospho-STAT1 and STAT2 indicates that the canonical ISGF3 complex is responsible for our observed induction of ISG15 in Δp53-Ras-shARF cells. To emphasize these findings, we overexpressed wild type STAT1 in Δp53 or Δp53-Ras cells and demonstrated by western blot analysis that overexpressed STAT1 alone was not able to increase ISG15 protein levels (Supplementary Fig. 3A) and was not sufficient for increased proliferation (cell growth and foci assays) in cells that retained ARF expression (Supplementary Fig. 3B-C). In particular, STAT1 was phosphorylated on Y701 in Δp53-Ras cells measured by western blot (Supplementary Fig. 3A), indicating that activation of STAT1 at this residue was not sufficient to drive ISG15 expression in cells expressing high levels of ARF.

**Fig. 3.**
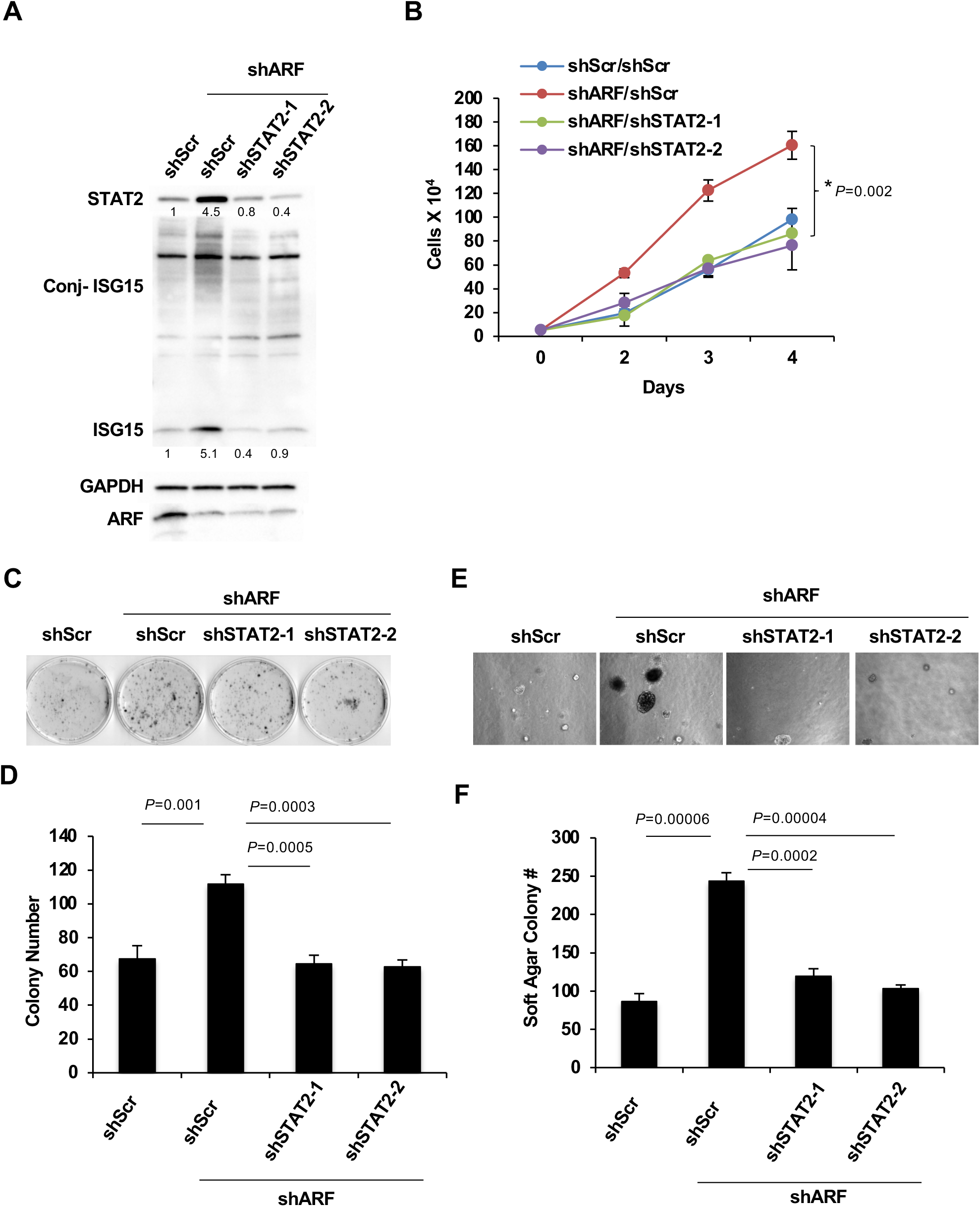
STAT2 is required for increased proliferation and transformation of Δp53-Ras- shARF MEFs. **A** Immunoblot analysis of cell lysates from Δp53-Ras-shScr or –shARF MEFs infected with two specific shRNAs targeting STAT2. Fold levels below each blot are relative to shScr control. Proteins analyzed are provided next to each blot. **B** Four-day proliferation assay performed with cells described in (A) with indicated *P* values for significant difference only. \**P*=0.0002 (purple or green compared to red). **C** Representative image of foci assay performed with cells described in (A). **D** Quantification of three independent measurements is shown with indicated *P* values. Error bars represent standard deviation (SD) of n=3. **E** Representative image of cells described in (A) growing in soft agar. **F** Macroscopic colonies were quantified (below) with indicated *P* values. Error bars represent standard deviation (SD) of n=3.

### Selective JAK inhibitors, ruxolitinib and baricitinib, decrease the proliferation of Δp53-Ras- shARF cells

Since the phosphorylation of STAT1-Y701 was critical for the increased proliferation and transformation of Δp53-Ras-shARF cells, we hypothesized that known JAK inhibitors, ruxolitinib and baricitinib, would mimic JAK1 knockdown through their effect on phospho-STAT1[47]. Treatment of Δp53-Ras-shARF MEFs with 1uM ruxolitinib or baricitinib decreased phospho- STAT1 levels and prevented the induction of ISG15 protein expression (Fig. 4A). Furthermore, Δp53-Ras-shARF MEFs that were treated with 1uM ruxolitinib or 1uM baricitinib exhibited decreased proliferation rates (Fig. 4B). This reduction was likely due to reduced proliferation, as we did not observe an increase in cleaved PARP, an indicator of apoptosis, after treatment with ruxolitinib or baricitinib (Supplemental Fig. 4). However, growth rates for Δp53-Ras-shScr MEFs were not altered after ruxolitinib or baricitinib treatment (Fig. 4B), indicating their complete lack of dependence on the type I interferon pathway.

**Fig. 4.**
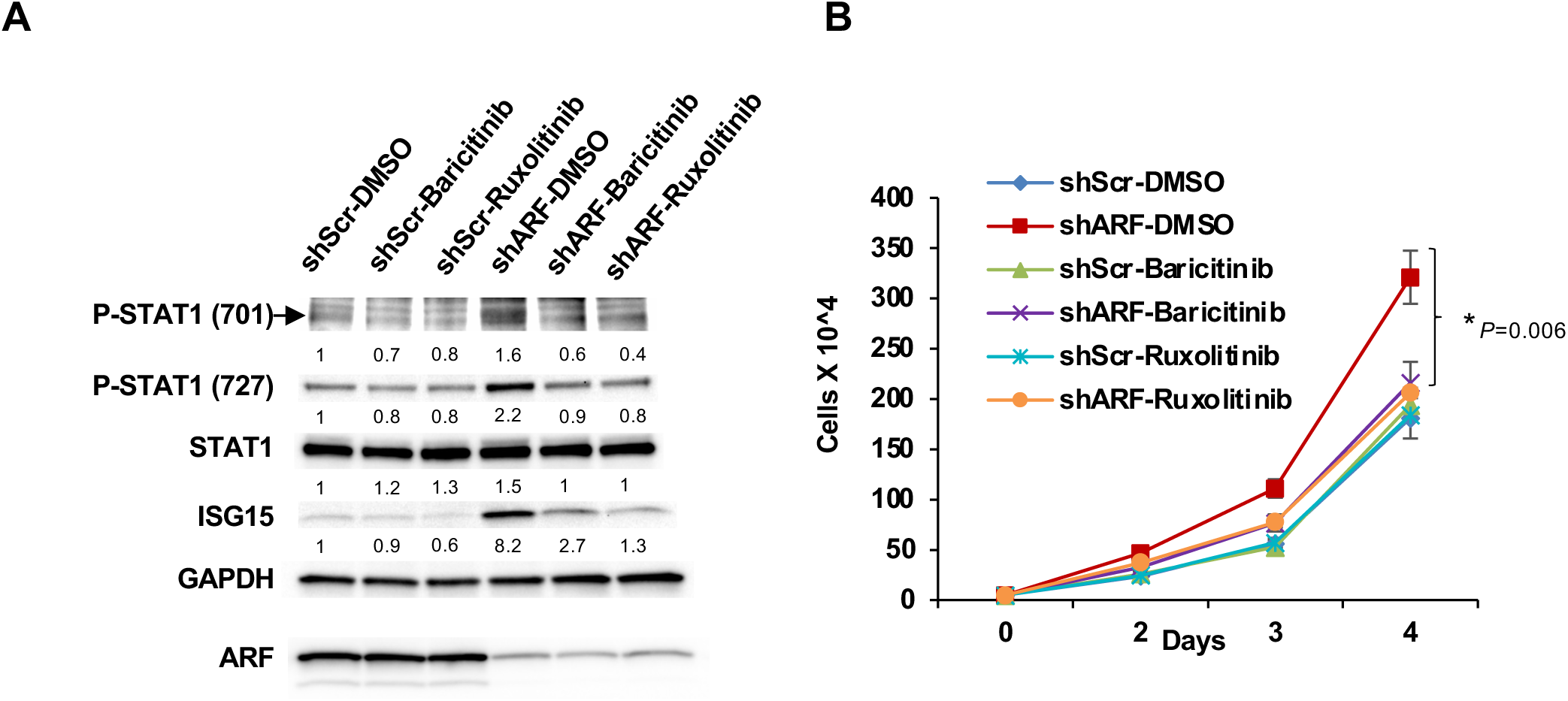
Δp53-Ras-shARF cells are sensitive to select JAK1 inhibitors. **A** Immunoblot analysis of cell lysates from Δp53-Ras-shScr or –shARF treated with vehicle control (DMSO), 1μM baricitinib, or 1μM ruxolitinib for 24 hrs. Proteins analyzed are provided next to each blot. Fold levels below each blot are relative to shScr control. **B** Four-day proliferation assay performed with cells described in (A) with indicated *P* values for significant difference only. \**P*=0.006 (purple or orange compared to red).

To better define the effects of JAK1inhibition on the signaling and proliferation of each MEF genotype, we treated MEFs with increasing doses of ruxolitinib. Δp53-Ras-shARF MEFs exhibited significant elevation of pSTAT1(701), total STAT1 and conjugated ISG15 protein levels compared to Δp53-Ras-shScr cells (Fig. 5A, B & D). Although 0.01μM ruxolitinib significantly reduced pSTAT1(701), total STAT1 and conjugated ISG15, these reduced protein levels were still much higher than in control cells (Fig. 5A, B &D). This was consistent with our proliferation findings, where 0.01μM did not significantly affect overall cell proliferation (Fig. 5C). However, treatment with 10μM ruxolitinib reduced pSTAT1(701), total STAT1, and conjugated ISG15 protein levels below control cells and resulted in attenuated cell proliferation (Fig. 5), suggesting that a threshold of pSTAT1, total STAT1, and conjugated ISG15 is required to maintain the increased proliferation seen in Δp53-Ras-shARF MEFs.

**Fig. 5.**
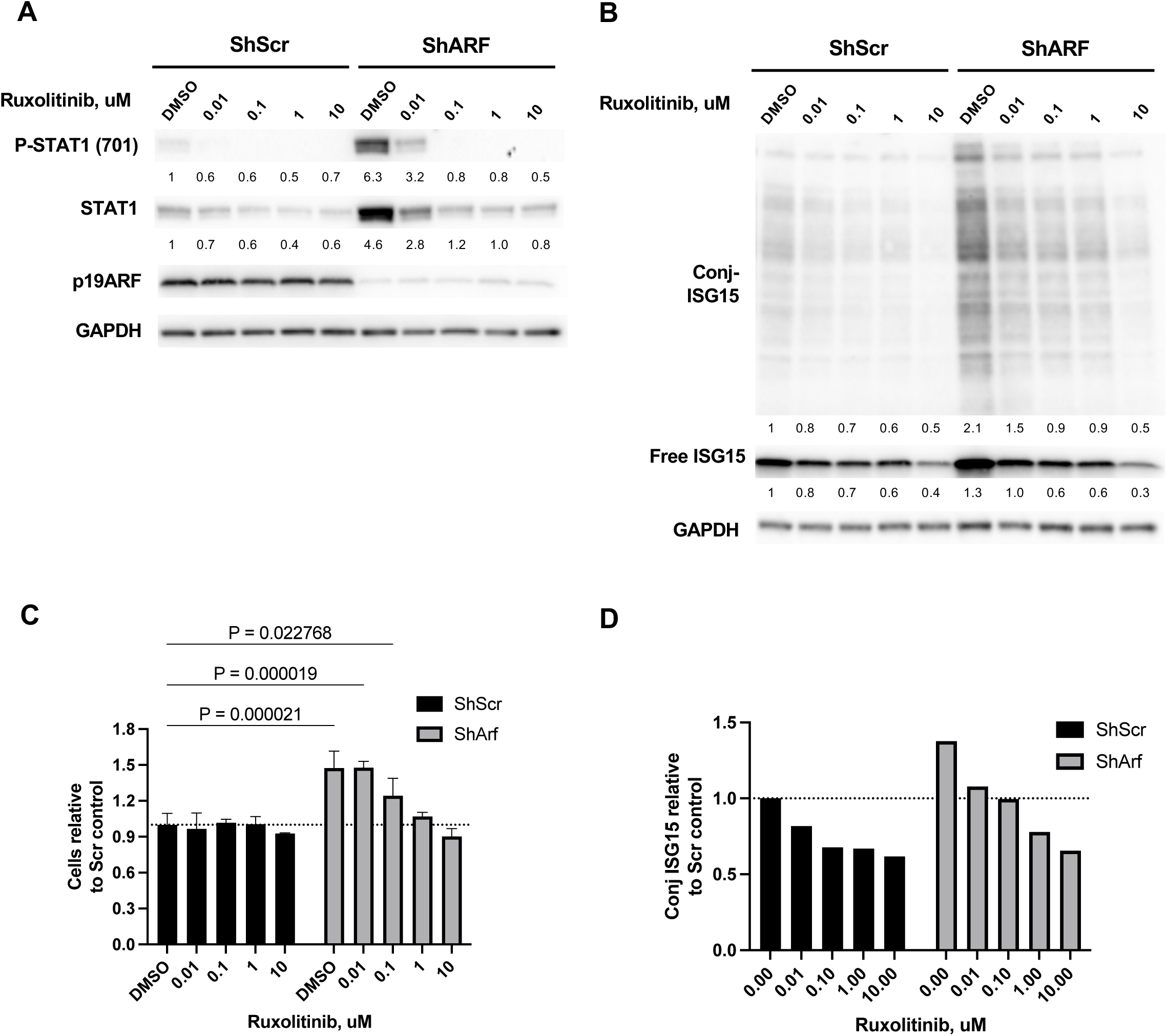
Dose response of Δp53-Ras-shARF cells to Ruxolitinib. **A** Immunoblot analysis of cell lysates from Δp53-Ras-shScr or –shARF treated with vehicle control (DMSO) or increasing concentrations of ruxolitinib for 24 hrs. Proteins analyzed are provided next to each blot. Fold levels below each blot are relative to shScr control. **B** Immunoblot analysis of free and conjugated ISG15 from Δp53-Ras-shScr or –shARF treated with vehicle control (DMSO) or increasing concentrations of ruxolitinib for 24 hrs. Proteins analyzed are provided next to each blot. Fold levels below each blot are relative to shScr control. **C** Δp53-Ras-shScr or –shARF treated with vehicle control (DMSO) or increasing concentrations of ruxolitinib for 24 hrs were counted and normalized to DMSO-shSCR cells where the dotted line=1.0. **D** immunoblot from (B) was analyzed for overall conjugated ISG15 where the dotted line=1.0.

### Loss of functional p53 and ARF is associated with poorer overall survival and activation of ISGs in multiple cancers

Our previous work has highlighted an important growth suppressive function of ARF in the context of p53 loss [18, 19]. Herein, we have now shown that these two tumor suppressors cooperate to inhibit cellular hyperproliferation driven by type I IFN production and JAK/STAT signaling. To determine to translational relevance of these findings, we explored human tumor data from The Cancer Genome Atlas (TCGA). We hypothesized that when p53 mutations and *CDKN2A* loss co-occur, these tumors would be more aggressive and express markers of elevated ISG signaling.

Using TCGA-PANCAN patient samples with both mutation and copy number data, we show that *CDKN2A* copy number loss on top of a p53 mutant background led to a significant reduction in overall survival (HR = 0.6691, 95% CI = 0.5765 to 0.7765, n=2080) compared to patients with p53 mutations alone (Fig 6A). Consistent with our mouse cell hyperproliferation data, patients with both *CDKN2A* loss and p53 mutation exhibited a significant increase in Ki-67 mRNA expression compared to all other genotypes (Fig. 6B). Next, we established that higher IFNβ, STAT1, and ISG15 mRNA expression together is pro-tumorigenic in patients and associated with reduced survival (HR = 0.8, 95% CI = 0.7221 to 0.8864, n=5682) (Fig. 6C). Breaking the expression of this signature down by TCGA study showed a diverse pattern among cancer types (Fig. 6D), suggestive of a tissue specific role for type I IFNs in cancer. To expand these results, we next stratified TCGA-PANCAN patients based on a low or high ISG score using ISG’s that were upregulated in *ARF/p53*-deficient MEFs [18]. Patients with high ISG expression were associated with reduced survival (HR = 0.6369, 95% CI = 0.5873 to 0.6907, n=6251) (Fig. 6E). Independent loss of function *CDKN2A* or p53 associated with a significantly increased ISG signature while loss of both *CDKN2A* and p53 mutation trended higher but was not statistically significant (Fig. 6F).

**Fig. 6.**
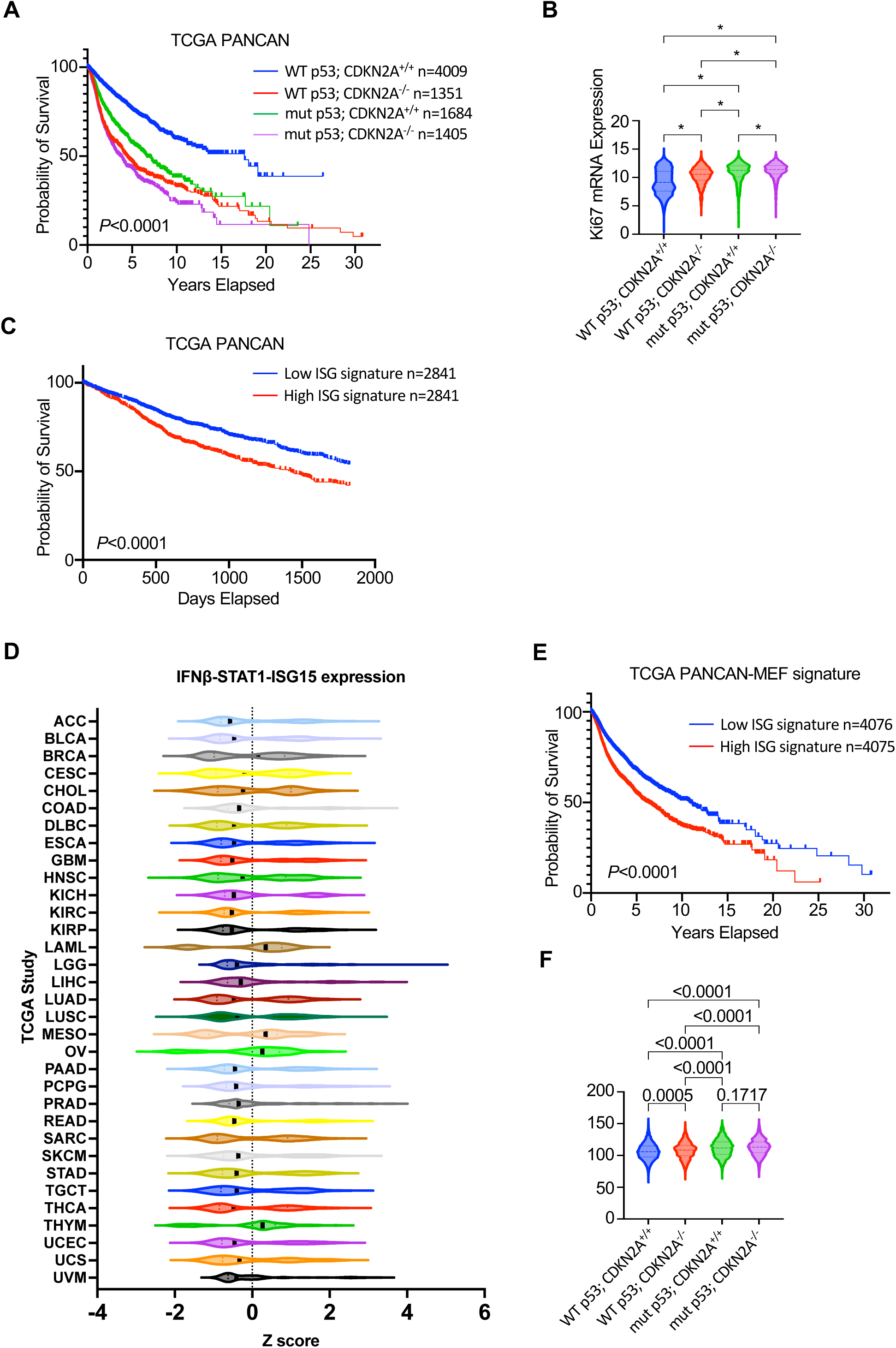
Impact of p53 and ARF loss in human cancer and Type I IFN pathway. **A** Overall survival curves were generated for pan-cancer patients exhibiting either wild type or mutant p53 with accompanying wild type CDKN2A or CDKN2A using TCGA data. **B** TCGA data from (A) were analyzed for Ki67 mRNA expression among the four indicated genotypes. *=*P*<0.001 **C** Overall survival curves were generated for pan-cancer patients exhibiting either a high (n=2841, red line) or low (n=2841, blue line) ISG gene expression signature. **D** TCGA data for the indicated human cancers were analyzed for IFNβ, STAT1, and ISG15 mRNA expression and plotted as a z score. **E** TCGA overall survival in all cancer data were analyzed for ISG expression (blue, ISG low and red, ISG high) based on the ISG gene signature observed in Δp53-Ras-shARF MEFs [18] **F** The indicated genotypes from all human cancers were analyzed for increased Δp53-Ras-shARF MEF ISG signature. *P* values are provided for significant difference only in comparing each genotype.

We hypothesized that the diversity of the PANCAN dataset may skew some of the findings based on inherent differences between cancer types. Based on previous murine data, loss of both *ARF* and *p53* in mice gives rise to several epithelial based tumors not seen in *p53*-null mice [19]. Therefore, we selected several common solid tumor types of epithelial origin to analyze the IFN pathway. We preformed gene set enrichment analysis (GSEA) between p53 mutated and p53 mutant/*CDKN2A* copy number loss samples within TCGA-BRCA (breast cancer), TCGA-LUAD (lung adenocarcinoma), TCGA-SKCM (melanoma), TCGA-PRAD (prostate adenocarcinoma), and TCGA-UCEC (uterine carcinoma/endometrial cancer). Remarkably, we found that the hallmark gene set for interferon alpha response, which captures genes for both IFNα and IFNβ responses, was significantly enriched with p53 mutant/*CDKN2A* loss among all these cancer types (Fig. 7A). We also uncovered a variety of gene sets related to cell cycle regulation and proliferation enriched in the p53 mutant*/CDKN2A* groups (Supplementary Excel Table1). The overlap between significantly enriched gene sets between cancer types is depicted in Fig. 7A. A closer analysis of the individual genes in the IFNα response gene set indicated genes that were shared among the five cancers to varying degrees (Fig 6B, Supplementary Fig. 5, Supplementary Excel Table2).

**Fig. 7.**
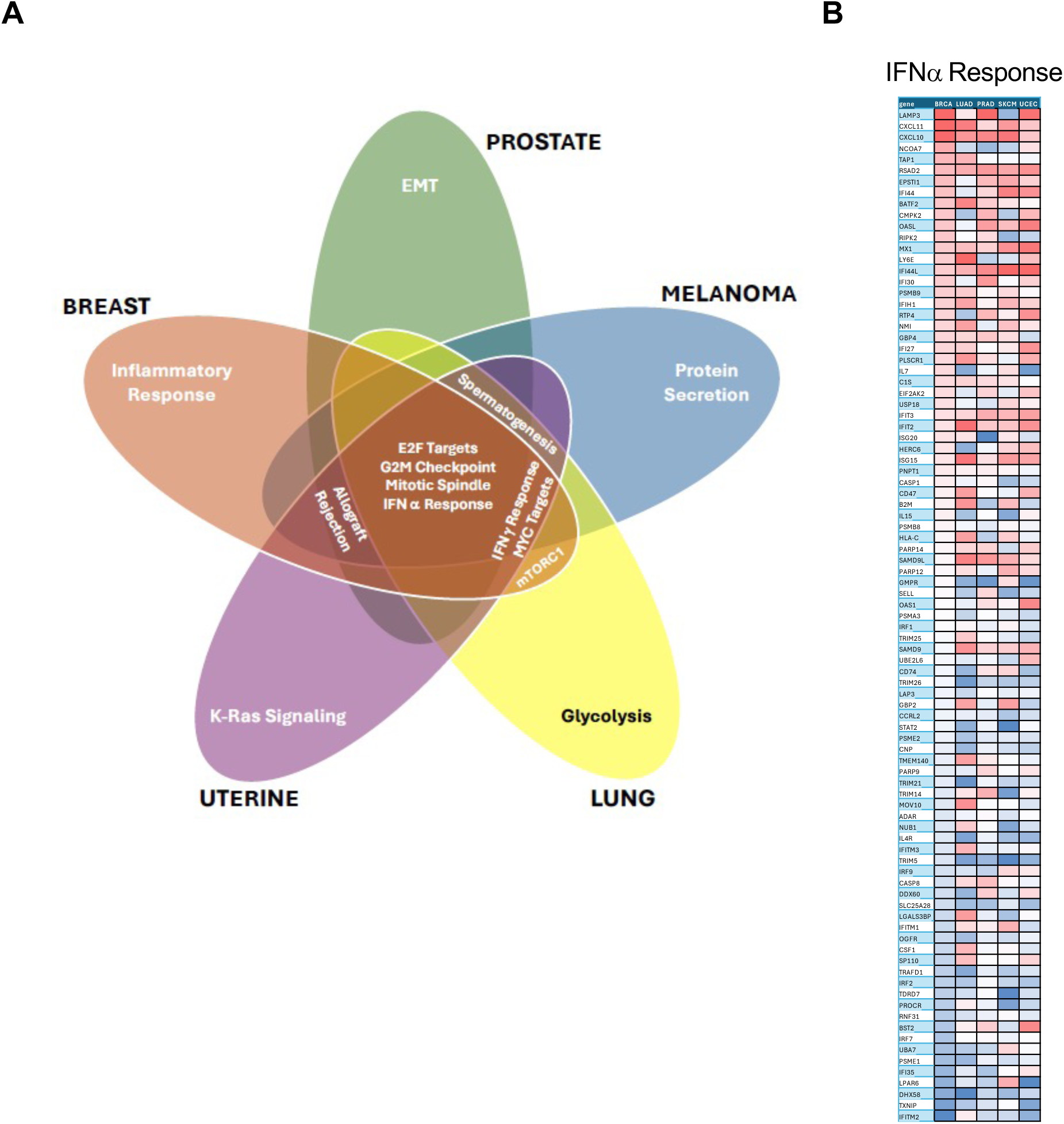
Shared pathways between p53 mutant breast, prostate, melanoma, lung, and uterine cancers. RNA sequencing data were analyzed from TCGA datasets for wild type versus mutant p53. A gene set enrichment analysis showed common pathways between multiple cancers. **A** Significant common pathways were plotted on a Venn diagram for breast (orange), lung adenocarcinoma (yellow), melanoma (blue), prostate (green), and uterine (purple) cancers. Overlap between cancers is shown and pathways denoted. **B** Genes corresponding to the IFNα response pathway were analyzed for changes in expression for each of the five indicated cancers. Expression changes between wild type and mutant p53 are denoted by an artificial color gradient of red (higher in mutant p53 samples) and blue (higher in wild type p53 samples).

### Shared pathways between p53 mutant breast, prostate, melanoma, lung, and uterine cancers

Having shown the importance of increased type I IFN signaling in the maintenance of accelerated proliferation in transformed mouse cells, we sought to extend these findings to human cancers. We initially performed an evaluation of the RNA sequencing data deposited into the TCGA for various human cancers. When available, we compared mRNA expression between wild type and mutant p53 samples within each cancer type. A gene set enrichment analysis (GSEA) of this comparison identified pathways that were increased in mutant p53 patient samples (Supplementary Excel Table 1). GSEA clustered five solid tumors together: breast, lung adenocarcinoma, melanoma, prostate, and uterine. These solid tumors had four pathways that were commonly elevated in the presence of mutant *p53*. mRNAs within the E2F targets, the G2M checkpoint, and the mitotic spindle pathways were elevated in each of the five solid cancers (Fig. 7A). This is consistent with the role of p53 in regulating each of these pathways and with that regulation being severely abrogated by the expression of the mutant p53 protein [48]. In particular, the type I IFN response was significantly induced in each of the five solid cancers harboring mutant *p53* (Fig. 7A). Other immune response pathways were shared among at least four of the five tumors. These included allograft rejection (breast, melanoma, prostate, and uterine) and IFNγ response (breast, lung, melanoma, and uterine). Breast cancer had three immune pathways in its top upregulated GSEA pathways, making it stand out among all solid tumors (Fig. 7A & Supplementary Excel Table 1). A heat map of mRNA expression levels in mutant versus wild type p53 samples indicated shared heightened expression of several IFNα response genes (Fig. 7B, Supplementary Fig. 5), underscoring the potential importance of this pathway in mediating the pathobiology of these tumors.

### Breast and lung cancers exhibit elevated ISG15 and sensitivity to selective JAK1 inhibitors

To test the idea that elevated type I IFN signaling might be at least partially responsible for the proliferation of cancer cells harboring mutant p53 alleles, we focused on breast and lung cancers. Consistent with our earlier findings in mice [18] and MEFs (see Figs. 2-4), breast cancers exhibiting concomitant mutations in *p53* and *CDKN2A* exhibited a significantly worse overall survival (Fig. 8A). To investigate this *in vitro*, we selected the 4T1 mouse mammary tumor cell line. 4T1 cells are null for *p53* and have deleted the *CDKN2A* locus (Fig. 8B). As a result, these cells show increased phosphorylation of STAT1(701), tremendous levels of ISG15 protein, and no p19ARF protein expression (Fig. 8B). Treatment of these cells with 10μM ruxolitinib resulted in a significantly reduced short-term proliferation (Fig. 8C) and long-term proliferation (Fig. 8D) and reduction in ISG15 expression (Supplementary Fig. 6A), demonstrating a role for JAK1 activity in the proliferation of these murine breast cancer cells. Human lung adenocarcinoma also showed a significantly decreased overall survival in patients with concomitantly mutated *p53* and *CDKN2A* that trend towards a co-mutation (*P*<0.001; Log odds ratio of 0.540; Fig. 8E). Next, we analyzed 24-lung adenocarcinomas for expression of the ARF and ISG15 protein by immunohistochemical analysis. Seventeen of the samples expressed mutant p53 along with low or undetectable p14ARF protein, representing 71% of the cases (Fig. 8F). Notably, all 17 of these tumors also exhibited high ISG15 protein staining compared to the 7 samples that expressed high p14ARF (Fig. 8F). We identified established lung cancer cell lines that expressed mutant p53 with high levels of p14ARF and undetectable IFNβ protein (HCC827) or mutant p53 with undetectable p14ARF and high levels of IFNβ protein (H838) (Fig. 8G). Treatment of these cells with 1μM baricitinib resulted in an attenuation of phosphorylated STAT3 (Supplemental Fig. 6B). This treatment also significantly reduced short-term (Supplemental Fig. 7A) and long-term proliferation of H838 but not HCC827 cells (Fig. 8H), underscoring the relationship between p14ARF, IFN signaling levels, and proliferation. Treatment with 10μM baricitinib nearly abolished all long-term proliferation in H838 cells, while HCC827 cells remained completely unaffected by even this high dose of baricitinib (Supplemental Fig. 7, C-D).

**Fig. 8.**
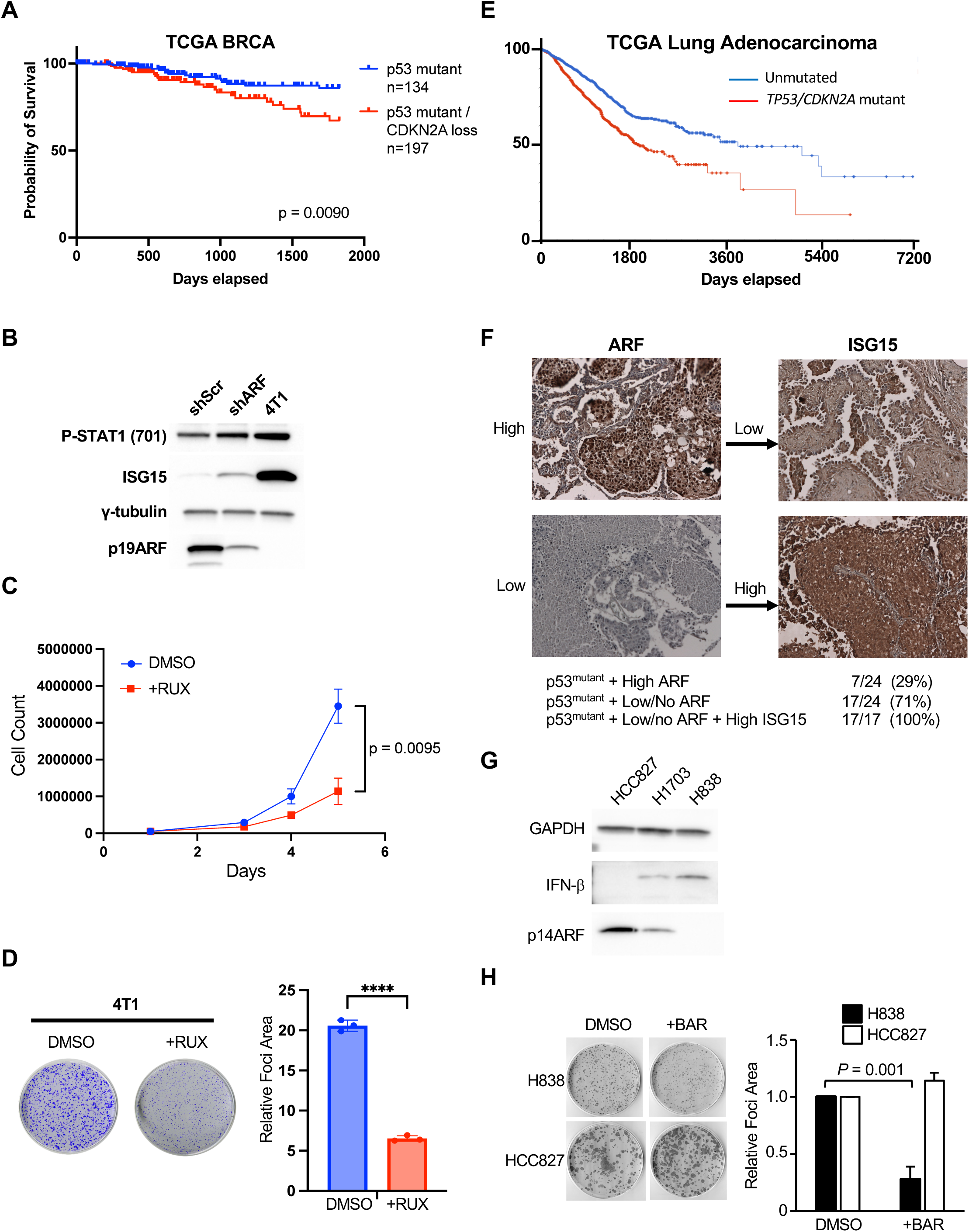
***p53* and *CDKN2A* status affects overall survival of human breast and lung cancer patients and cells from these cancers are sensitive to JAK1 inhibition. A** TCGA data from breast cancer patients was analyzed for overall survival in p53 mutant (134 cases) and p53 mutant with CDKN2A loss (197 cases). *P*=0.009. **B** Proteins isolated from mouse 4T1 breast cancer cells were separated by SDS-PAGE and immunoblotted with the indicated antibodies and compared to Δp53-Ras-shScr or –shARF cells. **C** Five-day proliferation assay performed with cells described in (B). **D** Representative image of foci assay performed with cells described in (B). Quantification of three independent measurements is shown with ****=*P*<0.001. Error bars represent standard deviation (SD) of n=3. **E** TCGA data from non-small cell lung cancer patients was analyzed for overall survival in p53 mutant (1446 cases) and p53 mutant with CDKN2A loss (1013 cases). *P*=2.32e-11. **F** Lung adenocarcinoma tissues were immunostained with antibodies recognizing p14ARF and ISG15. High and low expression for each protein is depicted from 24 total samples. **G** Proteins isolated from human lung cancer cells were separated by SDS-PAGE and immunoblotted with the indicated antibodies. **H** Representative image of foci assay performed with H838 and HCC827 cells that were treated with DMSO or 1μM baricitinib. Quantification of three biological replicates performed in triplicate is shown with indicated *P* values. Error bars represent standard deviation of n=3.

## DISCUSSION

In recent years, it has become evident that JAK/STAT signaling in response to type I IFN is a crucial component of tumor cells [48] and their environment [49]. Inflammation, for example, is an emerging hallmark of cancer cells, and STAT activation can induce inflammation and promote cell proliferation [50, 51]. Treatment of some types of cancers with type I IFN has been approved by the FDA, and many patients see a survival benefit [52]. Mounting evidence suggests STAT1 activation might promote the progression of certain tumors. STAT1 and many of its transcriptional targets have been found to be overexpressed in breast, lung, and cervical cancers [53–55]. While it could be argued that overexpression of STAT1 may be a “passenger” in human cancer, data from several human cancer studies indicates this may not be the case [56]. An interferon related DNA damage signature, which includes STAT1, predicts chemo and radiation therapy sensitivity in breast cancer patients [57]. Understanding which of these contexts STAT1 acts in a tumor-promoting fashion would provide a very accessible therapeutic target in these patients, as there are already STAT1 specific drugs as well as cell-permeable peptide inhibitors under investigation [58, 59]. Understanding the context in which type I IFN are produced in cancers would shed much light on the contrasting views of IFN, STAT1, and STAT2 in tumor biology [60–64].

In the setting of *ARF* and *p53* tumor suppressor deficiency, murine cells secrete IFNβ and signal through STAT1 to increase proliferation and transformation [18]. Stimulation of the canonical IFNβ pathway requires JAK1-mediated phosphorylation of STAT1 that in turn activates transcription of numerous genes including ISG15 [65]. Indeed, we have now shown that this canonical JAK1/STAT1 signaling pathway is required for the increased production of ISG15 in the absence of *p53* and *ARF*. Furthermore, phosphorylation of STAT1 on Y701 was critical for the increased proliferation and transformation of Δp53-Ras-shARF cells in stark contrast to a previous report that suggested a unphosphorylated STAT1 complex was responsible for chronic induction of ISG15 [60]. Other studies have suggested that phospho-STAT1 is pro-apoptotic thereby maintaining a tumor suppressive function, whereas unphosphorylated STAT1 is anti-apoptotic in sarcoma cells providing resistance to DNA damage [60, 66, 67]. However, our data contrasts these findings by identifying Y701 as a necessary site of JAK1-dependent phosphorylation in the absence of p53 and ARF, a genetic setting that could prove to be unique for the oncogenic function of phospho-STAT1.

We provide evidence that inhibition of JAK1 through shRNA-targeted knockdown or small molecules results in severe attenuation of proliferation only in *ARF/p53*-deficient cells, suggesting that this cellular context is uniquely vulnerable to JAK1 inhibition. Moreover, our data opens the potential use of JAK inhibitors such as ruxolitinib and baricitinib in tumor settings where p53 and ARF function are lost and increases in phospho-Y701 are evident. Ruxolitinib was recently used in a clinical trial for metastatic pancreatic cancer [68]; however, it was recently discontinued due to lack of sufficient efficacy. Our data argues that JAK1 inhibitors might be effective in decreasing the proliferation of tumors only in cancers where p53 and ARF are co-inactivated.

Building on this notion, our analysis of TCGA data show a clear indication of worse outcome when p53 and *CDKN2A* function are concomitantly lost. While *CDKN2A* encodes both the ARF and INK4A tumor suppressors, we have shown that INK4A does not function to suppress type I IFN production [18]. Moreover, the *ARF/p53*-double-null mice that exhibit elevated IFNβ production and STAT1 activity retain intact INK4A expression [19]. This is important as we cannot readily distinguish ARF and INK4A in TCGA datasets. TCGA data again were consistent with this idea, displaying worse prognosis in patients with an elevated ISG signature originally identified in *ARF/p53*-double-null mouse cells. Narrowing our focus in TCGA to identify specific tumor types with elevated ISG signatures, we were able to identify breast, uterine, prostate, lung, and melanoma tumors as having a gene set enrichment for a type I IFN response. This was striking given that the *ARF/p53*-double-null mouse model generates spontaneous epithelial-based carcinomas [19]. Breast cancer and lung cancer cell lines that exhibited alterations in both ARF and p53 and have an elevated IFN signature proved sensitive to selective JAK1 inhibitors, underscoring 1) that the genetic setting of ARF and p53 loss of function establishes the increased type I IFN response, and 2) that the increased type I IFN response sets the sensitivity to JAK1 inhibitors.

In summary, our study has identified a crucial role for type I IFN signaling in promoting tumor cell proliferation. We show that ARF and p53 loss of function results in increases in type I IFN production, JAK1 activity and STAT1/2 activity. These mouse findings translated to human cancers where we have now established a connection between ARF and p53 function with IFN signaling and ISG expression. Moreover, we discovered that when epithelial cancers lose ARF and p53 function, they engage in type I IFN production, thus uncovering their unique reliance on increased JAK1/STAT1/2 activities.

## MATERIALS and METHODS

### Cell culture

Primary MEFs were isolated as previously described [6]. Cells were maintained in Dulbecco’s modified Eagle’s medium supplemented with 10% fetal bovine serum, 2mM glutamine, 0.1 mM nonessential amino acids, and 1mM sodium pyruvate. Lung cancer cell lines H838, HCC827, and H1703 as well as 4T1 cells were obtained from ATCC (Manassas, VA) and cultured in RPMI 1640 with GLUTamax supplemented with 10% fetal bovine serum.

### Viral production and plasmids

MEFs were infected as previously described [18] to achieve Δp53-Ras-shScr and Δp53-Ras- shARF MEFs. For lentiviral production, 293Ts were co-transfected with Lipofectamine 2000 (Invitrogen), pCMV-VSVG, pCMVΔR8.2, and pLKO.1 puromycin (shRNA) or pLVX hygromycin (overexpression/rescue) constructs. 48 hours post transfection viral supernatants were harvested. Cells were infected with lentivirus supplemented with 8mg/ml protamine sulfate for 10- 16 hrs. Sequences for shRNAs can be found in supplemental experimental procedures.

Retroviral MSCV-STAT1 WT, Y701F, and S727A constructs were a gift from Dr. Robert Schreiber (Washington University of St. Louis). For knockdown rescue experiments wobble mutations in STAT1 retroviral plasmids were made using the following primers:

fwd: 5’AAAGTCATGGCTGCCGAAAATATCCCCGAGAATCCCCTGAAGTAT-3’

rev: 5’-ATACTTCAGGGGATTCTCGGGGATATTTTCGGCAGCCATGACTTT-3’

### Western blot analysis

Cells were lysed by sonication in EBC buffer (50mM Tris-Cl [pH 7.4], 120 mM NaCl, 0.5% NP- 40, 1mM EDTA) with PMSF and HALT Protease and Phosphatase Inhibitor cocktail (Thermo Scientific). 50ug of protein were run on SDS-polyacrylamide gels, transferred to PVDF (Millipore), and probed using the following antibodies: IFNβ, ADAR1, Υ-tubulin, p19, ISG15, (Santa Cruz Biotechnologies); MDA5, RIG-I, TRIM25, Jak1, JAK2, STAT1, p-STAT1Y701, p- STAT1S727 (Cell Signaling); STAT2 (Abcam); GAPDH (Bethyl Laboratories). The mouse ISG15 antibody was a gift from Dr. Deborah Lenschow or rabbit anti-ISG15 antibody (Invitrogen). HRP-conjugated secondary antibodies were from Jackson ImmunoResearch (West Grove, PA) Ruxolitinib and baricitinib were purchased from Selleck Chem and re-suspended in DMSO.

### Immunohistochemistry

Annotated lung cancer tissue arrays were obtained from US Biomax (Cat#T047). Staining was performed as previously described [18] using the Dako EnVision+ System-HRP (DAB) according to the manufacturer’s instructions. Rabbit anti-p14ARF (Bethyl) and mouse anti-ISG15 (Santa Cruz) were used at a 1:200 dilution. Quantification was performed by two separate individuals by blindly scoring staining intensity on a 0–3 scale, with 0 being no staining and 3 being strong widespread staining. A score of 0–1 was considered ‘low/no’ staining, and a score of 2–3 was considered ‘high’.

### Proliferation & Foci Assays

For proliferation assays, 5x10^4^ cells were plated in triplicate on six-well plates. Cells were harvested and counted with a hemacytometer on days 2, 3, and 4-post plating. For foci assays, 3x10^3^ cells were plated in triplicate on 10 cm dishes. 8-10 days (MEFs/4T1) or 21 days (lung cancer cells) post plating cells were fixed with 100% methanol and stained with Giemsa (Sigma Aldrich). Colonies were manually counted.

### Soft Agar Assay

For soft agar, 2.6x10^4^ cells were seeded in triplicate on 60-mm dishes as previously described[18]. Cells were layered with media/0.4% agar mix every 7 days. At 21 days, plates were examined under a microscope and colonies >0.5mm were manually counted.

### Apoptosis assay

Cells were stained with FITC-annexin V and propidium iodide using the Vybrant Apoptosis Assay Kit #3 (V13242; Molecular Probes/Invitrogen, Carlsbad, CA). For a positive control, cells were treated with etoposide (50uM) for 48 hours. Cells were analyzed by flow cytometry using a Becton Dickson FACSCalibur cell sorter with CELLQuest Pro (v5.2) analytical software.

### TCGA data analysis

TCGA survival and expression data was mined using USCS XenaBrowser (https://xenabrowser.net/heatmap/) [69]. For genotype-based survival analysis, TP53 status was determined using somatic mutation analysis from aggregated variable call format (VCF) data. CDKN2A status was determined using copy number data with a <-0.2 cutoff for loss. RNAseq gene expression data for *ifnb1, stat1,* and *isg15* was weighted, combined and Z transformed using XenaBrowser. The median signature expression was used as the cutoff for low/high survival groups. Statistical analysis preformed in GraphPad Prism v10. Significance (P < 0.05) for survival analysis was determined by Log-rank (Mantel-Cox) test.

Gene Set Enrichment Analysis (GSEA) was performed using BlitzGSEA (Ma’ayan Lab) through XenaBrowser using default parameters [70]. Genes within the HALLMARK_INTERFERON_ALPHA_RESPONSE gene set were displayed on a heatmap to show relative contribution to overall enrichment score.

### Statistical Analysis

Data were analyzed in GraphPad Prism (v10) using student t-test, one-way ANOVA with Dunnets’s correction for comparisons to a single control, or a simple linear regression model as indicated.

### shRNA Sequences

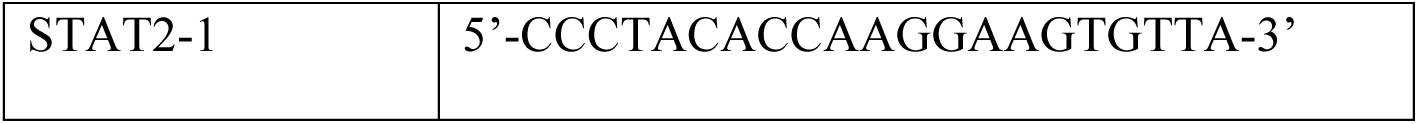
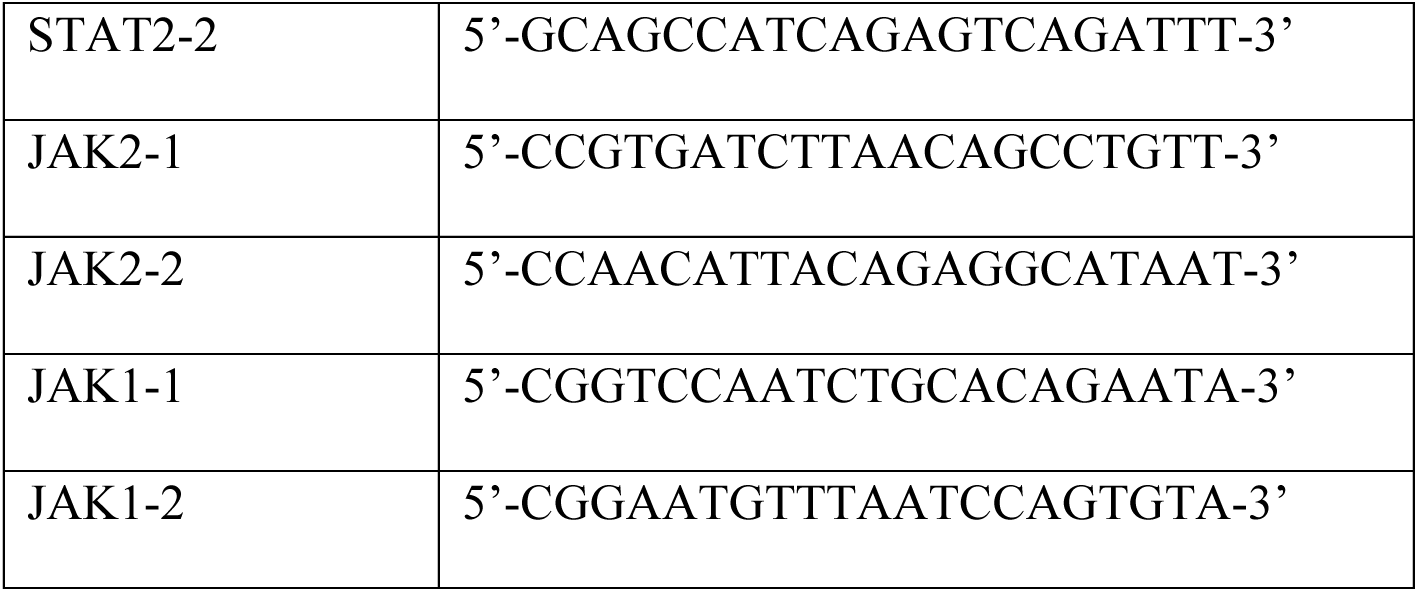

## Acknowledgements

We thank Deborah Lenschow for generously providing the ISG15 antibody and Robert Schreiber for providing the STAT1 expression plasmids. We also thank Jason Forys and other members of the Weber laboratory for their technical assistance. We are grateful to Katherine Weilbaecher, David DeNardo, Deborah Lenschow, and Susana Gonzalo for critical suggestions.

## Author contributions

C.E.K. conducted experiments, analyzed data, drafted, and edited the manuscript. A.M. conducted experiments, analyzed data, drafted, and edited the manuscript. A.O. conducted experiments, analyzed data, drafted, and edited the manuscript. L.B.M. conducted experiments, analyzed data, reviewed, and edited the manuscript. J.D.W. conceived the study, analyzed data, and wrote the manuscript.

## REFERENCES

1 Oren M. Regulation of the p53 tumor suppressor protein. J Biol Chem 1999; 274: 36031–36034.

2 Vousden KH. Activation of the p53 tumor suppressor protein. Biochim Biophys Acta 2002; 1602: 47–59.

3 Haupt Y, Maya R, Kazaz A, Oren M. Mdm2 promotes the rapid degradation of p53. Nature 1997; 387: 296–299.

4 Serrano M, Gomez-Lahoz E, DePinho RA, Beach D, Bar-Sagi D. Inhibition of ras-induced proliferation and cellular transformation by p16INK4. Science 1995; 267: 249–252.

5 Serrano M, Hannon GJ, Beach D. A new regulatory motif in cell-cycle control causing specific inhibition of cyclin D/CDK4. Nature 1993; 366: 704–707.

6 Kamijo T, Weber JD, Zambetti G, Zindy F, Roussel MF, Sherr CJ. Functional and physical interactions of the ARF tumor suppressor with p53 and Mdm2. Proc Natl Acad Sci U S A 1998; 95: 8292–8297.

7 Stott FJ, Bates S, James MC, McConnell BB, Starborg M, Brookes S et al. The alternative product from the human CDKN2A locus, p14(ARF), participates in a regulatory feedback loop with p53 and MDM2. EMBO J 1998; 17: 5001–5014.

8 Young NP, Jacks T. Tissue-specific p19Arf regulation dictates the response to oncogenic K-ras. Proc Natl Acad Sci U S A 2010; 107: 10184–10189.

9 Sreeramaneni R, Chaudhry A, McMahon M, Sherr CJ, Inoue K. Ras-Raf-Arf signaling critically depends on the Dmp1 transcription factor. Mol Cell Biol 2005; 25: 220–232.

10 Zindy F, Eischen CM, Randle DH, Kamijo T, Cleveland JL, Sherr CJ et al. Myc signaling via the ARF tumor suppressor regulates p53-dependent apoptosis and immortalization. Genes Dev 1998; 12: 2424–2433.

11 Eischen CM, Weber JD, Roussel MF, Sherr CJ, Cleveland JL. Disruption of the ARF- Mdm2-p53 tumor suppressor pathway in Myc-induced lymphomagenesis. Genes Dev 1999; 13: 2658–2669.

12 Zindy F, Williams RT, Baudino TA, Rehg JE, Skapek SX, Cleveland JL et al. Arf tumor suppressor promoter monitors latent oncogenic signals in vivo. Proc Natl Acad Sci U S A 2003; 100: 15930–15935.

13 Pomerantz J, Schreiber-Agus N, Liegeois NJ, Silverman A, Alland L, Chin L et al. The Ink4a tumor suppressor gene product, p19Arf, interacts with MDM2 and neutralizes MDM2’s inhibition of p53. Cell 1998; 92: 713–723.

14 Weber JD, Kuo ML, Bothner B, DiGiammarino EL, Kriwacki RW, Roussel MF et al. Cooperative signals governing ARF-mdm2 interaction and nucleolar localization of the complex. Mol Cell Biol 2000; 20: 2517–2528.

15 Weber JD, Taylor LJ, Roussel MF, Sherr CJ, Bar-Sagi D. Nucleolar Arf sequesters Mdm2 and activates p53. Nat Cell Biol 1999; 1: 20–26.

16 Lohrum MA, Ashcroft M, Kubbutat MH, Vousden KH. Identification of a cryptic nucleolar- localization signal in MDM2. Nat Cell Biol 2000; 2: 179–181.

17 Riley T, Sontag E, Chen P, Levine A. Transcriptional control of human p53-regulated genes. Nat Rev Mol Cell Biol 2008; 9: 402–412.

18 Forys JT, Kuzmicki CE, Saporita AJ, Winkeler CL, Maggi LB, Jr., Weber JD. ARF and p53 coordinate tumor suppression of an oncogenic IFN-beta-STAT1-ISG15 signaling axis. Cell Rep 2014; 7: 514–526.

19 Weber JD, Jeffers JR, Rehg JE, Randle DH, Lozano G, Roussel MF et al. p53- independent functions of the p19(ARF) tumor suppressor. Genes Dev 2000; 14: 2358–2365.

20 Brady SN, Yu Y, Maggi LB, Jr., Weber JD. ARF impedes NPM/B23 shuttling in an Mdm2- sensitive tumor suppressor pathway. Mol Cell Biol 2004; 24: 9327–9338.

21 Sugimoto M, Kuo ML, Roussel MF, Sherr CJ. Nucleolar Arf tumor suppressor inhibits ribosomal RNA processing. Mol Cell 2003; 11: 415–424.

22 Bertwistle D, Sugimoto M, Sherr CJ. Physical and functional interactions of the Arf tumor suppressor protein with nucleophosmin/B23. Mol Cell Biol 2004; 24: 985–996.

23 Itahana K, Bhat KP, Jin A, Itahana Y, Hawke D, Kobayashi R et al. Tumor suppressor ARF degrades B23, a nucleolar protein involved in ribosome biogenesis and cell proliferation. Mol Cell 2003; 12: 1151–1164.

24 Tago K, Chiocca S, Sherr CJ. Sumoylation induced by the Arf tumor suppressor: a p53- independent function. Proc Natl Acad Sci U S A 2005; 102: 7689–7694.

25 Saporita AJ, Chang HC, Winkeler CL, Apicelli AJ, Kladney RD, Wang J et al. RNA helicase DDX5 is a p53-independent target of ARF that participates in ribosome biogenesis. Cancer Res 2011; 71: 6708–6717.

26 Lessard F, Morin F, Ivanchuk S, Langlois F, Stefanovsky V, Rutka J et al. The ARF tumor suppressor controls ribosome biogenesis by regulating the RNA polymerase I transcription factor TTF-I. Mol Cell 2010; 38: 539–550.

27 Rozenblum E, Schutte M, Goggins M, Hahn SA, Panzer S, Zahurak M et al. Tumor-suppressive pathways in pancreatic carcinoma. Cancer Res 1997; 57: 1731–1734.

28 Sanchez-Cespedes M, Reed AL, Buta M, Wu L, Westra WH, Herman JG et al. Inactivation of the INK4A/ARF locus frequently coexists with TP53 mutations in non-small cell lung cancer. Oncogene 1999; 18: 5843–5849.

29 Cerami E, Gao J, Dogrusoz U, Gross BE, Sumer SO, Aksoy BA et al. The cBio cancer genomics portal: an open platform for exploring multidimensional cancer genomics data. Cancer Discov 2012; 2: 401–404.

30 Gao J, Aksoy BA, Dogrusoz U, Dresdner G, Gross B, Sumer SO et al. Integrative analysis of complex cancer genomics and clinical profiles using the cBioPortal. Sci Signal 2013; 6: pl1.

31 Marie I, Durbin JE, Levy DE. Differential viral induction of distinct interferon-alpha genes by positive feedback through interferon regulatory factor-7. EMBO J 1998; 17: 6660–6669.

32 Colamonici OR, Uyttendaele H, Domanski P, Yan H, Krolewski JJ. p135tyk2, an interferon-alpha-activated tyrosine kinase, is physically associated with an interferon- alpha receptor. J Biol Chem 1994; 269: 3518–3522.

33 Novick D, Cohen B, Rubinstein M. The human interferon alpha/beta receptor: characterization and molecular cloning. Cell 1994; 77: 391–400.

34 Fu XY, Schindler C, Improta T, Aebersold R, Darnell JE, Jr. The proteins of ISGF-3, the interferon alpha-induced transcriptional activator, define a gene family involved in signal transduction. Proc Natl Acad Sci U S A 1992; 89: 7840–7843.

35 Hu X, Herrero C, Li WP, Antoniv TT, Falck-Pedersen E, Koch AE et al. Sensitization of IFN-gamma Jak-STAT signaling during macrophage activation. Nat Immunol 2002; 3: 859–866.

36 Morales DJ, Lenschow DJ. The antiviral activities of ISG15. J Mol Biol 2013; 425: 4995–5008.

37 Platanias LC. Mechanisms of type-I- and type-II-interferon-mediated signalling. Nat Rev Immunol 2005; 5: 375–386.

38 Chan SR, Vermi W, Luo J, Lucini L, Rickert C, Fowler AM et al. STAT1-deficient mice spontaneously develop estrogen receptor alpha-positive luminal mammary carcinomas. Breast Cancer Res 2012; 14: R16.

39 Klover PJ, Muller WJ, Robinson GW, Pfeiffer RM, Yamaji D, Hennighausen L. Loss of STAT1 from mouse mammary epithelium results in an increased Neu-induced tumor burden. Neoplasia 2010; 12: 899–905.

40 Hix LM, Karavitis J, Khan MW, Shi YH, Khazaie K, Zhang M. Tumor STAT1 transcription factor activity enhances breast tumor growth and immune suppression mediated by myeloid-derived suppressor cells. J Biol Chem 2013; 288: 11676–11688.

41 Chen RH, Du Y, Han P, Wang HB, Liang FY, Feng GK et al. ISG15 predicts poor prognosis and promotes cancer stem cell phenotype in nasopharyngeal carcinoma. Oncotarget 2016; 7: 16910–16922.

42 Song L, Rawal B, Nemeth JA, Haura EB. JAK1 activates STAT3 activity in non-small-cell lung cancer cells and IL-6 neutralizing antibodies can suppress JAK1-STAT3 signaling. Mol Cancer Ther 2011; 10: 481–494.

43 Wong GS, Lee JS, Park YY, Klein-Szanto AJ, Waldron TJ, Cukierman E et al. Periostin cooperates with mutant p53 to mediate invasion through the induction of STAT1 signaling in the esophageal tumor microenvironment. Oncogenesis 2013; 2: e59.

44 Schindler C, Shuai K, Prezioso VR, Darnell JE, Jr. Pillars article: Interferon-dependent tyrosine phosphorylation of a latent cytoplasmic transcription factor. Science. 1992. 257: 809-813. J Immunol 2011; 187: 5489–5494.

45 Shuai K, Schindler C, Prezioso VR, Darnell JE, Jr. Activation of transcription by IFN-gamma: tyrosine phosphorylation of a 91-kD DNA binding protein. Science 1992; 258: 1808–1812.

46 Shuai K, Stark GR, Kerr IM, Darnell JE, Jr. A single phosphotyrosine residue of Stat91 required for gene activation by interferon-gamma. Science 1993; 261: 1744–1746.

47 Roskoski R, Jr. Janus kinase (JAK) inhibitors in the treatment of inflammatory and neoplastic diseases. Pharmacol Res 2016; 111: 784–803.

48 Cheon H, Wang Y, Wightman SM, Jackson MW, Stark GR. How cancer cells make and respond to interferon-I. Trends Cancer 2023; 9: 83–92.

49 Gonzalez-Navajas JM, Lee J, David M, Raz E. Immunomodulatory functions of type I interferons. *Nature reviews Immunology* (Research Support, N.I.H., Extramural Research Support, Non-U.S. Gov’t Review) 2012; 12: 125–135.

50 Hanahan D, Weinberg RA. Hallmarks of cancer: the next generation. Cell (Research Support, N.I.H, Extramural Review) 2011; 144: 646–674.

51 Yu H, Pardoll D, Jove R. STATs in cancer inflammation and immunity: a leading role for STAT3. Nature reviews Cancer (Review) 2009; 9: 798–809.

52 Dunn GP, Koebel CM, Schreiber RD. Interferons, immunity and cancer immunoediting. Nature reviews Immunology (Research Support, N.I.H., Extramural Research Support, Non-U.S. Gov’t Review) 2006; 6: 836–848.

53 Perou CM, Jeffrey SS, van de Rijn M, Rees CA, Eisen MB, Ross DT et al. Distinctive gene expression patterns in human mammary epithelial cells and breast cancers. *Proceedings of the National Academy of Sciences of the United States of America* (Research Support, Non-U.S. Gov’t Research Support, U.S. Gov’t, P.H.S.) 1999; 96: 9212–9217.

54 Rajkumar T, Sabitha K, Vijayalakshmi N, Shirley S, Bose MV, Gopal G et al. Identification and validation of genes involved in cervical tumourigenesis. *BMC cancer* (Research Support, Non-U.S. Gov’t Validation Studies) 2011; 11: 80.

55 Govindan R, Ding L, Griffith M, Subramanian J, Dees ND, Kanchi KL et al. Genomic landscape of non-small cell lung cancer in smokers and never-smokers. *Cell* (Research Support, N.I.H., Extramural Research Support, Non-U.S. Gov’t) 2012; 150: 1121–1134.

56 Khodarev NN, Roizman B, Weichselbaum RR. Molecular pathways: interferon/stat1 pathway: role in the tumor resistance to genotoxic stress and aggressive growth. *Clinical cancer research : an official journal of the American Association for Cancer Research* (Research Support, N.I.H., Extramural Research Support, Non-U.S. Gov’t) 2012; 18: 3015–3021.

57 Weichselbaum RR, Ishwaran H, Yoon T, Nuyten DS, Baker SW, Khodarev N et al. An interferon-related gene signature for DNA damage resistance is a predictive marker for chemotherapy and radiation for breast cancer. Proceedings of the National Academy of Sciences of the United States of America (Research Support, N.I.H., Extramural Validation Studies) 2008; 105: 18490–18495.

58 Wang H, Yang Y, Sharma N, Tarasova NI, Timofeeva OA, Winkler-Pickett RT et al. STAT1 activation regulates proliferation and differentiation of renal progenitors. Cellular signalling (Research Support, N.I.H., Intramural) 2010; 22: 1717–1726.

59 Sikorski K, Czerwoniec A, Bujnicki JM, Wesoly J, Bluyssen HA. STAT1 as a novel therapeutical target in pro-atherogenic signal integration of IFNgamma, TLR4 and IL-6 in vascular disease. Cytokine & growth factor reviews (Research Support, Non-U.S. Gov’t Review) 2011; 22: 211–219.

60 Cheon H, Holvey-Bates EG, Schoggins JW, Forster S, Hertzog P, Imanaka N et al. IFNbeta-dependent increases in STAT1, STAT2, and IRF9 mediate resistance to viruses and DNA damage. EMBO J 2013; 32: 2751–2763.

61 Weichselbaum RR, Ishwaran H, Yoon T, Nuyten DS, Baker SW, Khodarev N et al. An interferon-related gene signature for DNA damage resistance is a predictive marker for chemotherapy and radiation for breast cancer. Proc Natl Acad Sci U S A 2008; 105: 18490–18495.

62 Burnette BC, Liang H, Lee Y, Chlewicki L, Khodarev NN, Weichselbaum RR et al. The efficacy of radiotherapy relies upon induction of type i interferon-dependent innate and adaptive immunity. Cancer Res 2011; 71: 2488–2496.

63 Doherty MR, Cheon H, Junk DJ, Vinayak S, Varadan V, Telli ML et al. Interferon-beta represses cancer stem cell properties in triple-negative breast cancer. Proc Natl Acad Sci U S A 2017; 114: 13792–13797.

64 Khodarev NN, Beckett M, Labay E, Darga T, Roizman B, Weichselbaum RR. STAT1 is overexpressed in tumors selected for radioresistance and confers protection from radiation in transduced sensitive cells. Proc Natl Acad Sci U S A 2004; 101: 1714–1719.

65 Schindler C, Levy DE, Decker T. JAK-STAT signaling: from interferons to cytokines. J Biol Chem 2007; 282: 20059–20063.

66 Chin YE, Kitagawa M, Su WC, You ZH, Iwamoto Y, Fu XY. Cell growth arrest and induction of cyclin-dependent kinase inhibitor p21 WAF1/CIP1 mediated by STAT1. Science 1996; 272: 719–722.

67 Malilas W, Koh SS, Kim S, Srisuttee R, Cho IR, Moon J et al. Cancer upregulated gene 2, a novel oncogene, enhances migration and drug resistance of colon cancer cells via STAT1 activation. Int J Oncol 2013; 43: 1111–1116.

68 Hurwitz HI, Uppal N, Wagner SA, Bendell JC, Beck JT, Wade SM, 3rd et al. Randomized, Double-Blind, Phase II Study of Ruxolitinib or Placebo in Combination With Capecitabine in Patients With Metastatic Pancreatic Cancer for Whom Therapy With Gemcitabine Has Failed. J Clin Oncol 2015; 33: 4039–4047.

69 Goldman MJ, Craft B, Hastie M, Repecka K, McDade F, Kamath A et al. Visualizing and interpreting cancer genomics data via the Xena platform. Nat Biotechnol 2020; 38: 675–678.

70 Lachmann A, Xie Z, Ma’ayan A. blitzGSEA: efficient computation of gene set enrichment analysis through gamma distribution approximation. Bioinformatics 2022; 38: 2356–2357.

