## Supplemental Figures for "Elevated type I interferon signaling defines the proliferative advantage of ARF and p53 mutant tumor cells"

### Supplemental Figure 1

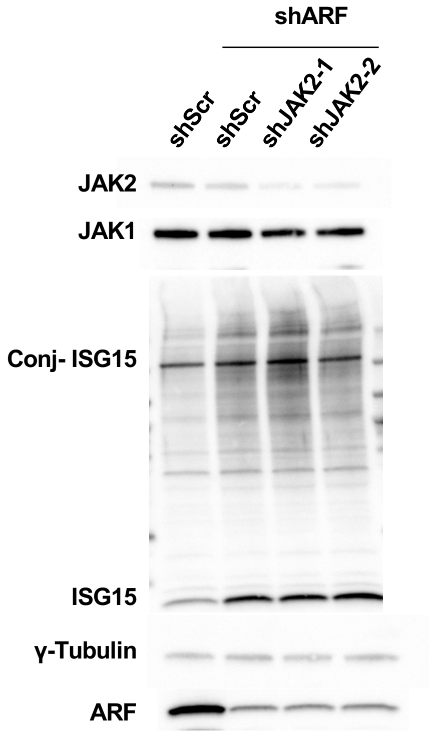

**Supplemental Figure 2**

**A**

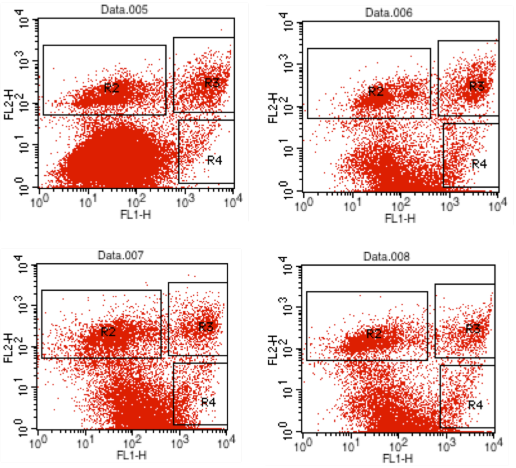


| Sample | % Cells Undergoing Apoptosis |
|---|---|
| shScr/shScr | 3.68 |
| shARF/shScr | 4.31 |
| shARF/shJAK1-1 | 5.33 |
| shARF/shJAK1-2 | 3.29 |

**B**

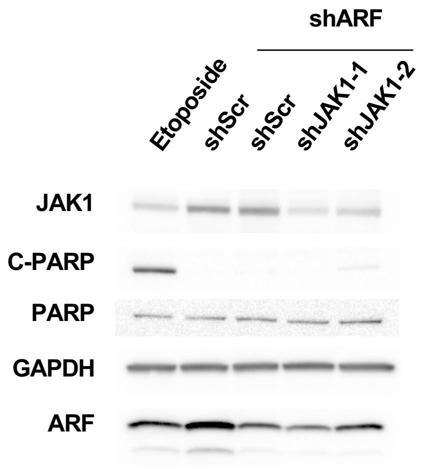

**Supplemental Figure 3**

**A**

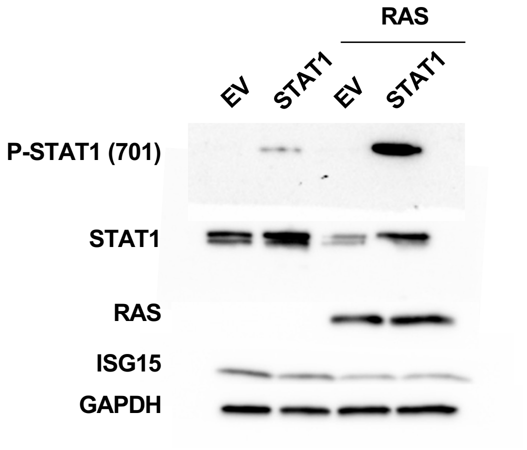


**B**

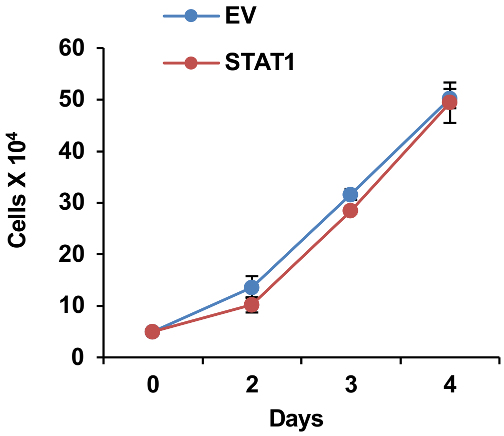


**C**

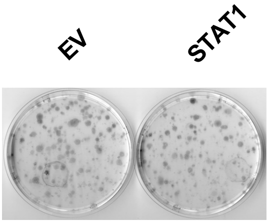

**Supplemental Figure 4**

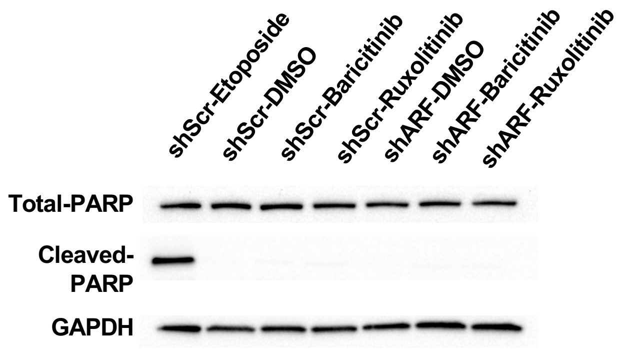

**Supplemental Figure 5**

BRCA

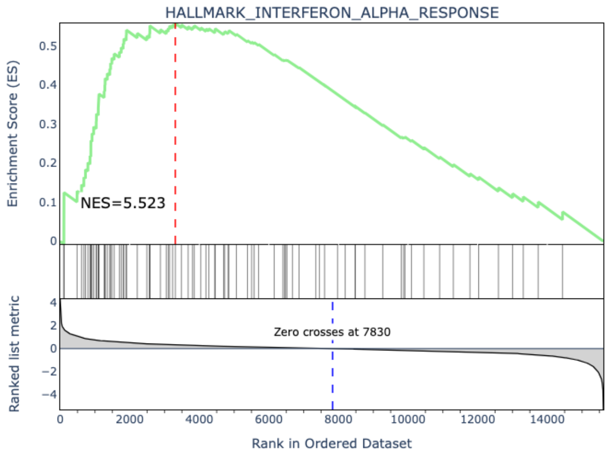


UCEC

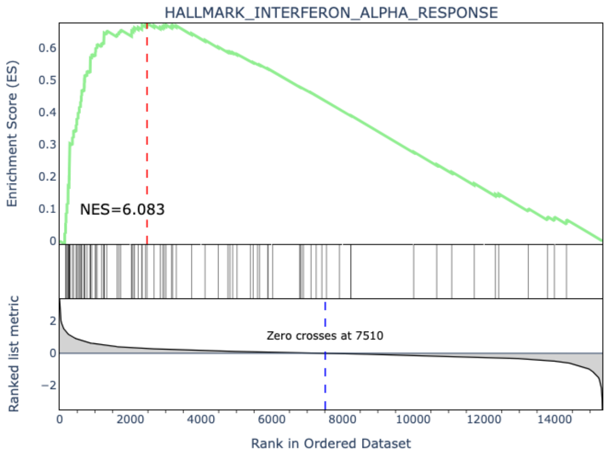


PRAD

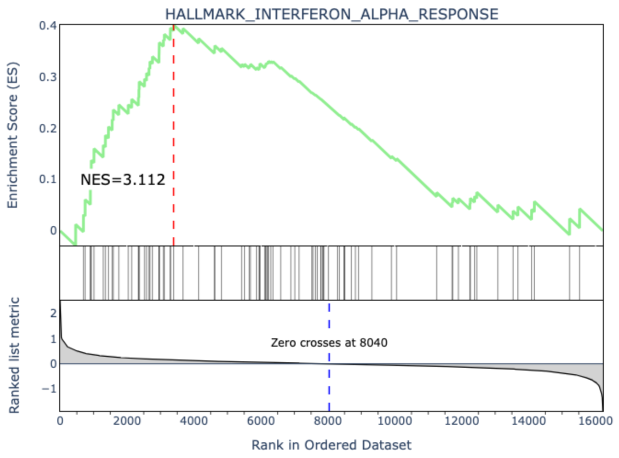


LUAD

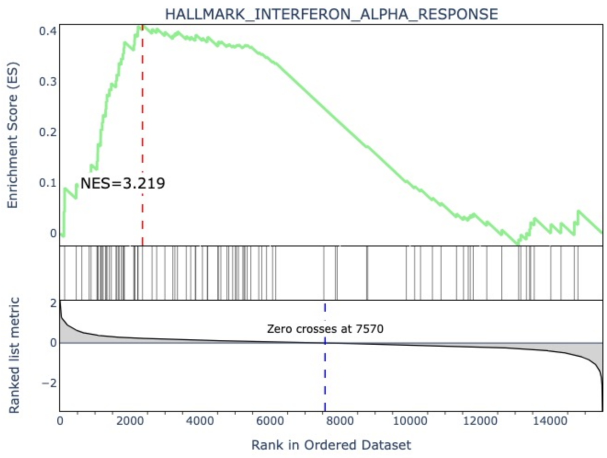


SKCM

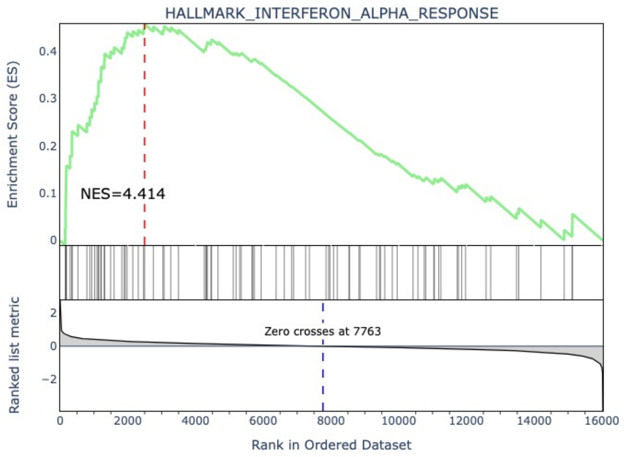

**Supplemental Figure 6**

**A**

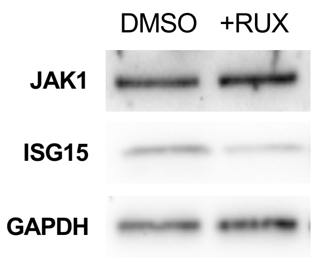


**B**

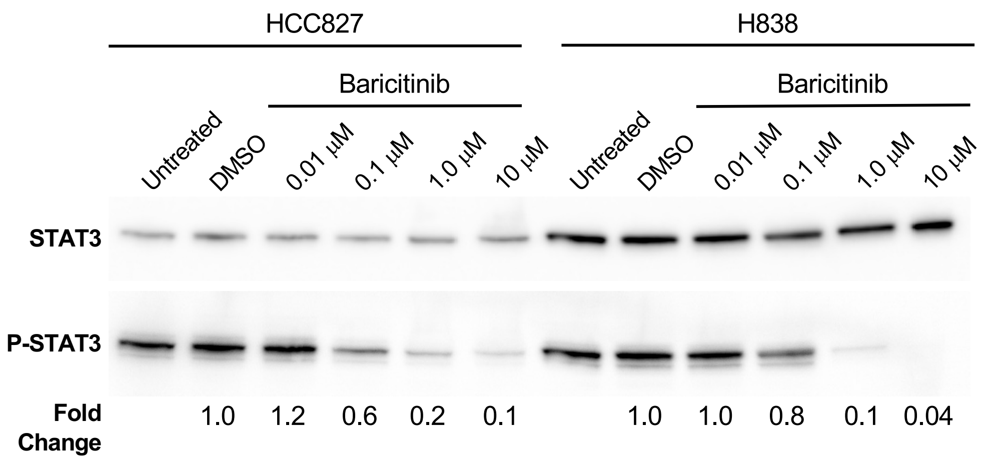

**Supplemental Figure 7**

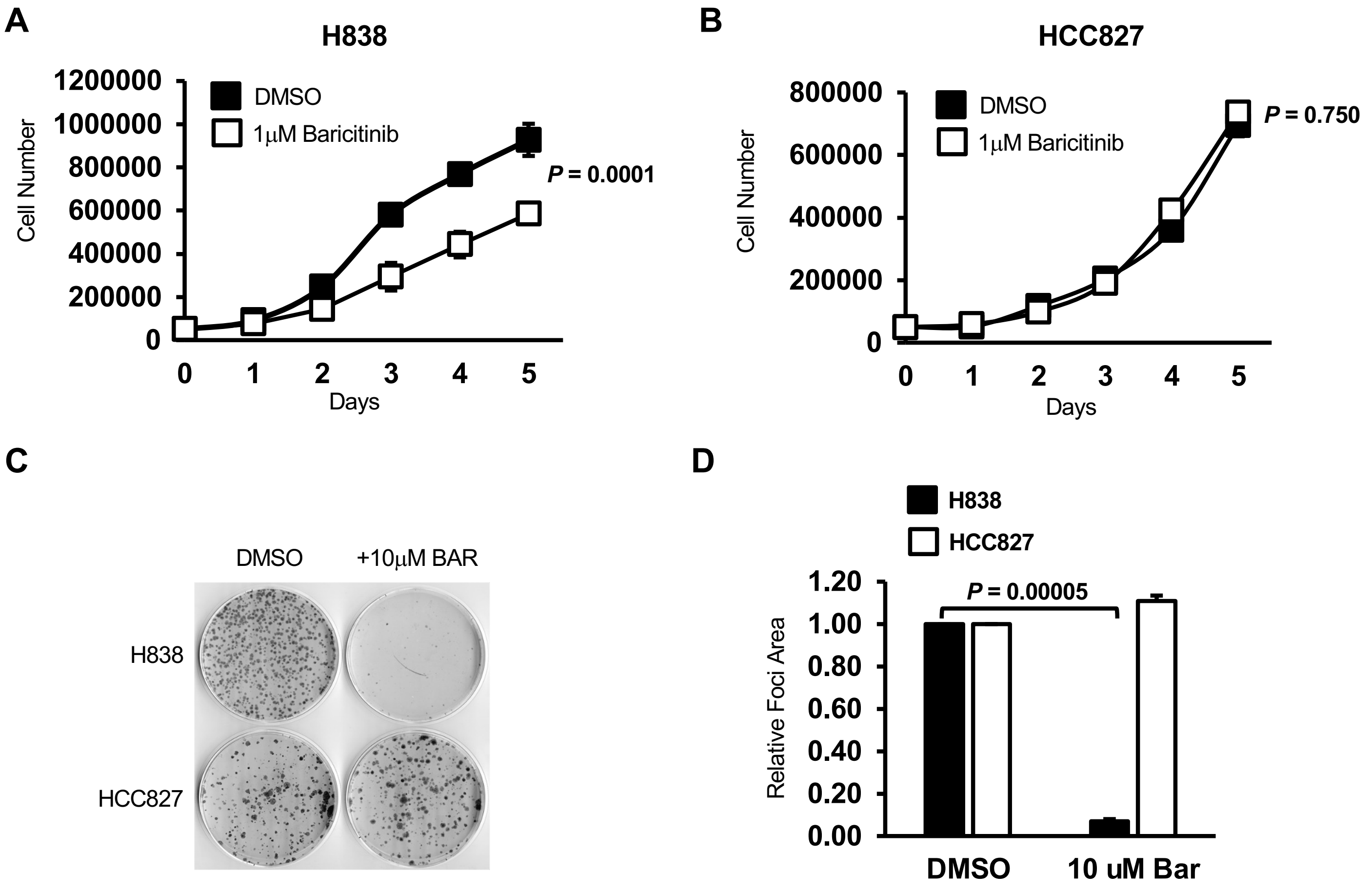
