## Supplemental Figure Legends for "Elevated type I interferon signaling defines the proliferative advantage of ARF and p53 mutant tumor cells"

**Supplemental Figure 1. Knockdown of JAK2 does not affect free and conjugated ISG15 in Δp53-Ras-shARF cells.** Immunoblot analysis of cell lysates from Δp53-Ras-shScr or –shARF MEFs infected with two specific shRNAs targeting JAK2 using the indicated antibodies.

**Supplemental Figure 2. Lack of apoptosis in Δp53-Ras-shARF with reduced JAK1 expression.** **A** Cells described in Figure 1a were stained with FITC-annexin V and propidium iodide and analyzed by flow cytometry for R3 (late apoptosis) + R4 (early apoptosis). **B** Immunoblot analysis of cell lysates from Δp53-Ras-shScr or –shARF MEFs infected with two specific shRNAs targeting JAK1 using the indicated antibodies.

**Supplemental Figure 3. Overexpression of STAT1 does not affect ISG15 expression or proliferation of Δp53-Ras cells. A** Immunoblot analysis of cell lysates from Δp53-shScr or –shARF, or Δp53-Ras-shScr or-shARF MEFs infected with a STAT1 expression construct. **B** Proliferation assay performed with cells described in (A). **C** Representative image of foci assay performed with cells described in (A).

**Supplemental Figure 4. Selective JAK1 inhibitors do not induce cleaved PARP.** Δp53-Ras-shScr or –shARF MEFs were treated with etoposide (50μM), DMSO, baricitinib (1μM), or ruxolitinib (1μM) and harvested protein lysates were immunoblotted using the indicated antibodies.

**Supplemental Figure 5. Gene set enrichment analysis of human cancers.** Genes enriched in the interferon alpha response pathway were examined for BRCA (breast cancer), UCEC (uterine carcinoma endometrial cancer), PRAD (prostate adenocarcinoma), LUAD (lung adenocarcinoma), and SKCM (skin cancer melanoma).

**Supplemental Figure 6. Selective JAK1 inhibitors reduce ISG15 in 4T1 cells and phosphorylated STAT3 in lung cancer cells.** **A** 4T1 cells were treated with DMSO or 1μM ruxolitinib (+Rux). Proteins were harvested 24 hrs later, separated by SDS-PAGE, and immunoblotted with the indicated antibodies. **B** H838 or HCC827 cells (1x10^5^ cells/3 mL RPMI) were left untreated or treated with the indicated concentrations of Barcitinib for 24 h. Cell lysates were analyzed for phosphorylated and total STAT3 by Western blot. Changes in Phosphorylated STAT3 relative to total STAT3 are shown. Western blot analysis is representative of 2 independent experiments.

**Supplemental Figure 7. Baricitinib reduces the proliferation of lung cancer cells**. **a** H838 (A) or HCC827 (B) lung cancer cells (5x10^4^/ 3mL DMEM) were treated with DMSO or 1 μM Barcitinib for the indicated times at 37°C. Cells were counted each day for Growth Curve analysis. Results are the average ± SEM of three independent experiments, A simple linear regression model was used to determine significant differences between the growth curves; *P*=0.0001 (A) and *P*=0.750 (B). (C) H838 cells (3,000 cells/3 mL RPMI) were treated with DMSO or 10 μM Barcitinib for 21 days at 37°C with media changes every 7 days. Cells were stained with Gimsea and foci area measured (D). Foci Area Analysis is the average ± SEM of three independent experiments with a one-way ANOVA with Dunnett’s post hoc correction; P=0.00005 shown for significant differences only. Images are representative of three independent experiments.
